# Bacterial mimicry of eukaryotic HECT ubiquitin ligation

**DOI:** 10.1101/2023.06.05.543783

**Authors:** Tyler G. Franklin, Peter S. Brzovic, Jonathan N. Pruneda

**Author notes:** Address correspondence to Jonathan N. Pruneda.

## Abstract

HECT E3 ubiquitin (Ub) ligases direct their modified substrates toward a range of cellular fates dictated by the specific form of monomeric or polymeric Ub (polyUb) signal that is attached. How polyUb specificity is achieved has been a longstanding mystery, despite extensive study ranging from yeast to human. Two outlying examples of bacterial “HECT-like” (bHECT) E3 ligases have been reported in the human pathogens Enterohemorrhagic *Escherichia coli* and *Salmonella* Typhimurium, but what parallels can be drawn to eukaryotic HECT (eHECT) mechanism and specificity had not been explored. Here, we expanded the bHECT family and identified catalytically active, *bona fide* examples in both human and plant pathogens. By determining structures for three bHECT complexes in their primed, Ub-loaded states, we resolved key details of the full bHECT Ub ligation mechanism. One structure provided the first glimpse of a HECT E3 ligase in the act of ligating polyUb, yielding a means to rewire the polyUb specificity of both bHECT and eHECT ligases. Through studying this evolutionarily distinct bHECT family, we have not only gained insight into the function of key bacterial virulence factors but also revealed fundamental principles underlying HECT-type Ub ligation.

## INTRODUCTION

Ubiquitination is a critical post-translational modification that regulates a gamut of cellular processes ranging from targeted protein degradation to signal transduction. The ubiquitination pathway requires orchestration of a ubiquitin (Ub)-activating E1, Ub-conjugating E2, and E3 Ub ligase to modify substrates^1^. A distinguishing feature of the Homologous to E6AP C-terminus (HECT) E3 ligases is their ability to directly influence the substrate’s cellular fate through formation of distinct polymeric Ub (polyUb) signals that recruit different cellular response factors^1–3^. For example, the founding member of the HECT family, E6AP, is specific for lysine (Lys or K)48-linked polyUb^4, 5^ and can target substrates for proteasomal degradation^6, 7^, while Rsp5 adds K63-linked polyUb onto its targets during endocytic processes^8, 9^. Mutations that disrupt these regulatory processes are frequently observed in cancers and neurodegenerative disorders, among other diseases, making them crucial research targets^10^. Despite significant effort, however, a clear picture for how HECT E3 ligases catalyze ubiquitination is lacking.

As an alternative approach to understanding the mechanism of Ub ligation in eukaryotic HECT E3 ligases (eHECTs), we turned to a family of related enzymes in bacteria. While the complete ubiquitination pathway is present only in eukaryotes, microbial pathogens secrete Ub-targeted effector proteins to dysregulate the host Ub system in ways that benefit invasion, persistence, and replication^11^. Several classes of these bacterial effector proteins can functionally mimic eukaryotic E3s and insert themselves into the host ubiquitination pathway, including bacterial U-box E3s that function similarly to eukaryotic RING/U-box E3s^12^, as well as the HECT-like effector proteins SopA from *Salmonella enterica* Typhimurium and NleL from Enterohemorrhagic *Escherichia coli* (EHEC)^13, 14^. Crystal structures of NleL and SopA revealed structurally distinct but topologically similar HECT domains, with an E2-binding N-lobe and catalytic C-lobe joined by a linker region^15^. Similar to eHECTs, the bacterial HECT-like E3 ligases (bHECTs) also feature HECT-like domains at the protein C-terminus, with substrate-binding regions located upstream that mediate interactions with host factors^15–17^. While extensive work has demonstrated how eHECTs interact with Ub, E2, and E2∼Ub during ligation, it remains largely unknown how bHECTs interact with Ub, or even E2∼Ub in the process of catalyzing ubiquitination^4, 18–33^.

Like many of their eukaryotic counterparts, bHECTs also assemble specific types of polyUb signals. Interestingly, the bHECT NleL robustly generates K6-linked polyUb as a ∼50:50 mixture with K48-linked polyUb, representing the most K6-specific ligase known to-date^14, 34^. A clear understanding for the role of NleL and the K6-linked polyUb signals it generates is lacking, though several reports would indicate a connection with actin pedestals formed by EHEC^16, 35^. Meanwhile, the only other reported bHECT, SopA, preferentially generates K48-linked polyUb and has been tied to the Ub-dependent degradation of its targeted host factors, TRIM56 and TRIM65^11, 17, 36^. How NleL and SopA are able to dictate their polyUb products, and whether any of these mechanisms also mimic those used by eHECTs, remains an open question. The generally accepted model of polyUb chain formation by HECT E3s involves simultaneous coordination of two Ub molecules: a donor Ub (Ub^D^) that is transiently bound to the active site cysteine (Cys) of the HECT C-lobe, and an acceptor Ub (Ub^A^) that is optimally oriented so that the correct Lys residue performs nucleophilic attack^37, 38^. Among eHECTs, this polyUb linkage specificity appears to be partially encoded in the very C-terminal residues of the C-lobe^18, 30^. Still, a mechanism for how HECT E3 ligases catalyze specific polyUb signals largely remains a mystery.

Here, to elucidate the mechanisms of Ub ligation, we first expanded the bHECT family to include additional validated examples from both human and plant pathogens. Crystal structures of three bHECTs – NleL, SopA, and VsHECT – bound to Ub^D^ at their active sites revealed key features of this catalytic intermediate. These structures, combined with NMR data, identified commonalities between bHECT- and eHECT-mediated Ub ligation. Crystal packing of the NleL-Ub^D^ structure revealed the acceptor site for K48-linked polyUb ligation, providing the first visualization of a HECT:Ub^A^ interface^1, 37^. By mutating this Ub^A^ interface, K48-linked polyUb ligation by bHECTs could be redirected to K6-linked polyUb. Illustrating the functional mimicry of eHECT ligases, insights from the NleL:Ub^A^ interface informed mutational analyses of the eHECT HUWE1 that redirected its specificity toward increased K6-linked polyUb ligation. Thus, despite considerable differences in sequence and structure, bHECTs follow many of the same underlying principles of Ub ligation as their eukaryotic counterparts.

## RESULTS

### Expansion of the bacterial HECT-like E3 Ub ligase family

Unlike other bacterial E3 ligase families that are widely distributed among human and plant pathogens^39–42^, the HECT-like E3 ligase family was restricted to only two reported examples^13, 14^. To better appreciate the mechanism of bHECT ligases, we first used sequence and structural homology to identify other potential family members in pathogenic bacteria (see **Methods**) (**Fig. 1A**). Candidate sequences were prioritized based on their similarity to the canonical features of HECT-like ligases, including 1) an aromatic residue in the putative N-lobe E2 interaction site, 2) a potential C-lobe catalytic Cys residue ∼30 amino acids upstream of the C-terminus, and 3) a linker region bridging the N- and C-lobes (**Fig. 1A**)^14, 15, 37^. Though it was not used as a selection criterion, many bHECT candidates also encoded an N-terminal β-helix domain that is likely involved in substrate recognition^17^. bHECT candidates were found in both human and plant pathogen genomes, with relatively low amino acid conservation across the bHECT domain as well as individual regions (**Fig. 1B, S1A-C**). We selected bHECT candidates from *Proteus vulgaris* (PvHECT), *Verrucomicrobia* spp. (VsHECT), *Erwinia amylorova* (EaHECT), and *Proteus stewartii* (PsHECT), for testing E3 ligase activity of recombinantly purified protein (**Table S1**). Ub ligase activity was first determined using gel-based readouts for PvHECT, PsHECT, and VsHECT, in addition to the known bHECTs NleL and SopA, all of which consumed monomeric Ub to produce free polyUb chains and/or bHECT auto-ubiquitination (**Fig. 1C**). Mutation of the predicted active site Cys ablated ligase activity in all the newly identified bHECTs (**Fig. 1A, C**). Time-dependent ligase activity was additionally observed using the fluorescence polarization (FP) method UbiReal, which we have previously used to monitor bHECT and eHECT ligation^43, 44^. To varying degrees, addition of PsHECT, EaHECT, PvHECT, and VsHECT all produced a rise in FP of TAMRA-labeled Ub over time, indicating the presence of ligase activity (**Fig. S1D**).

**Figure 1:**
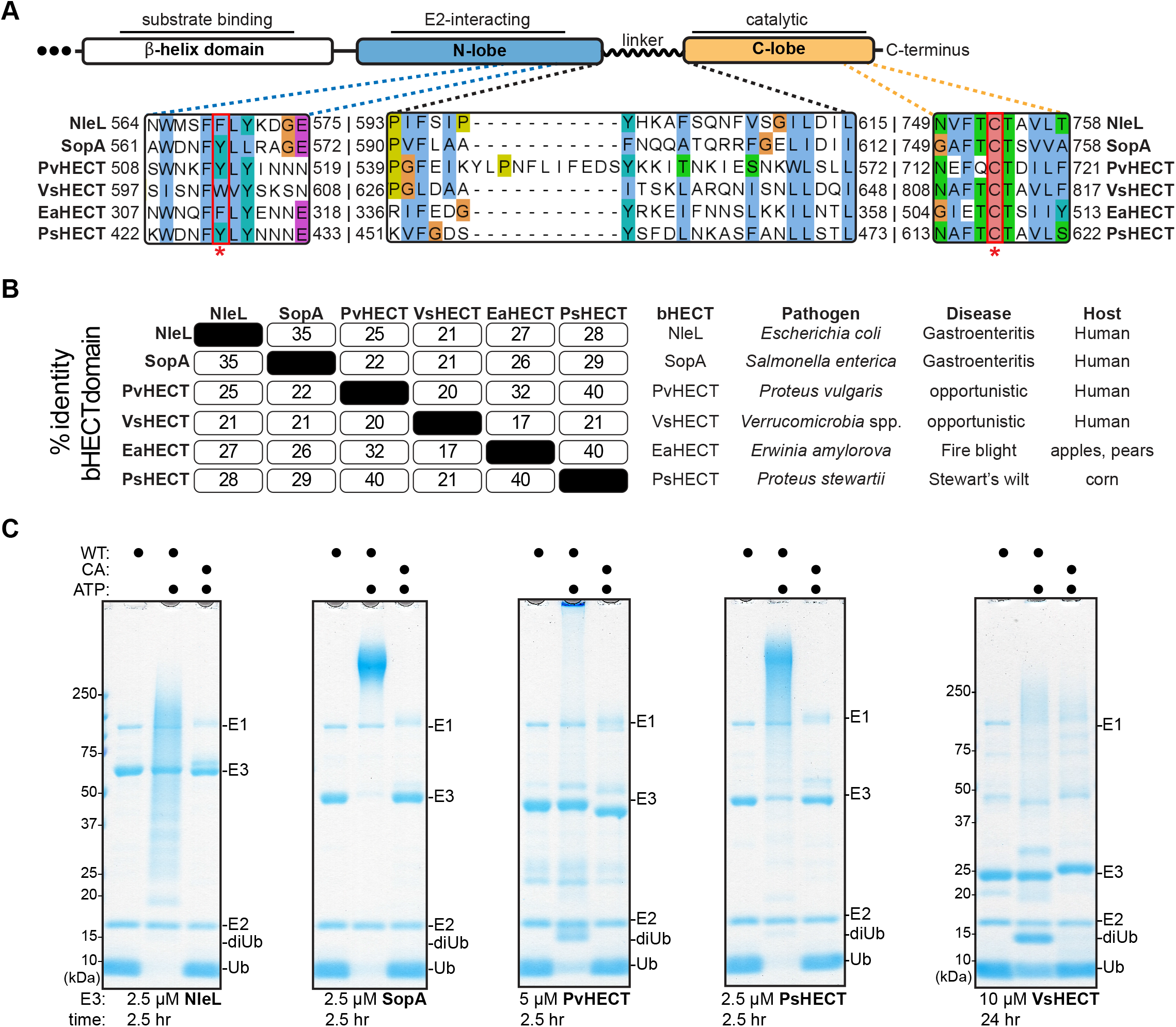
Discovery of an expanded bHECT family. A. Domain architecture of the HECT-like domain of bHECTs. Known critical regions, including the N-lobe aromatic residue, the C-lobe active site Cys, and the linker domain, are expanded to show sequence conservation at these sites. B. Percent sequence identity matrix for the entire HECT-like domain of the bHECTs, along with species of origin, presenting disease, and host. C. Gel-based Ub ligase assay for WT or the active site Cys mutant (CA) bHECTs. Reactions were initiated with ATP. bHECT concentrations are listed, and samples were taken at the indicated timepoints, quenched, and resolved by SDS-PAGE with Coomassie staining. See also Figure S1.

### Crystal structures reveal mechanisms of donor Ub coordination by bHECTs

Notably, the only soluble expression construct of VsHECT that we could obtain was the minimal C-lobe domain, yet weak ligase activity was still observed despite the lack of an E2-binding N-lobe (**Fig. 1C**). Ligase activity was also observed with minimal C-lobe constructs of NleL and SopA, though kinetics were reduced compared to the full bHECT domains (**Fig. 2A**). To further show that the C-lobe was the minimal catalytic region, we tested reactivity against the Ub-Propargylamide (PA) activity-based probe, which has previously been used to profile eHECTs and other Ub regulators^23, 45, 46^. For all bHECTs tested, we observed strong reactivity consistent with a single modification event of the active site cysteine (**Fig. 2B**). Notably, for eHECTs, reactivity with the Ub-PA probe is not observed in the absence of the N-lobe^23^. Thus, at least for bHECTs, the isolated C-lobe domain represents a minimal ligase module for studying Ub transfer events.

**Figure 2:**
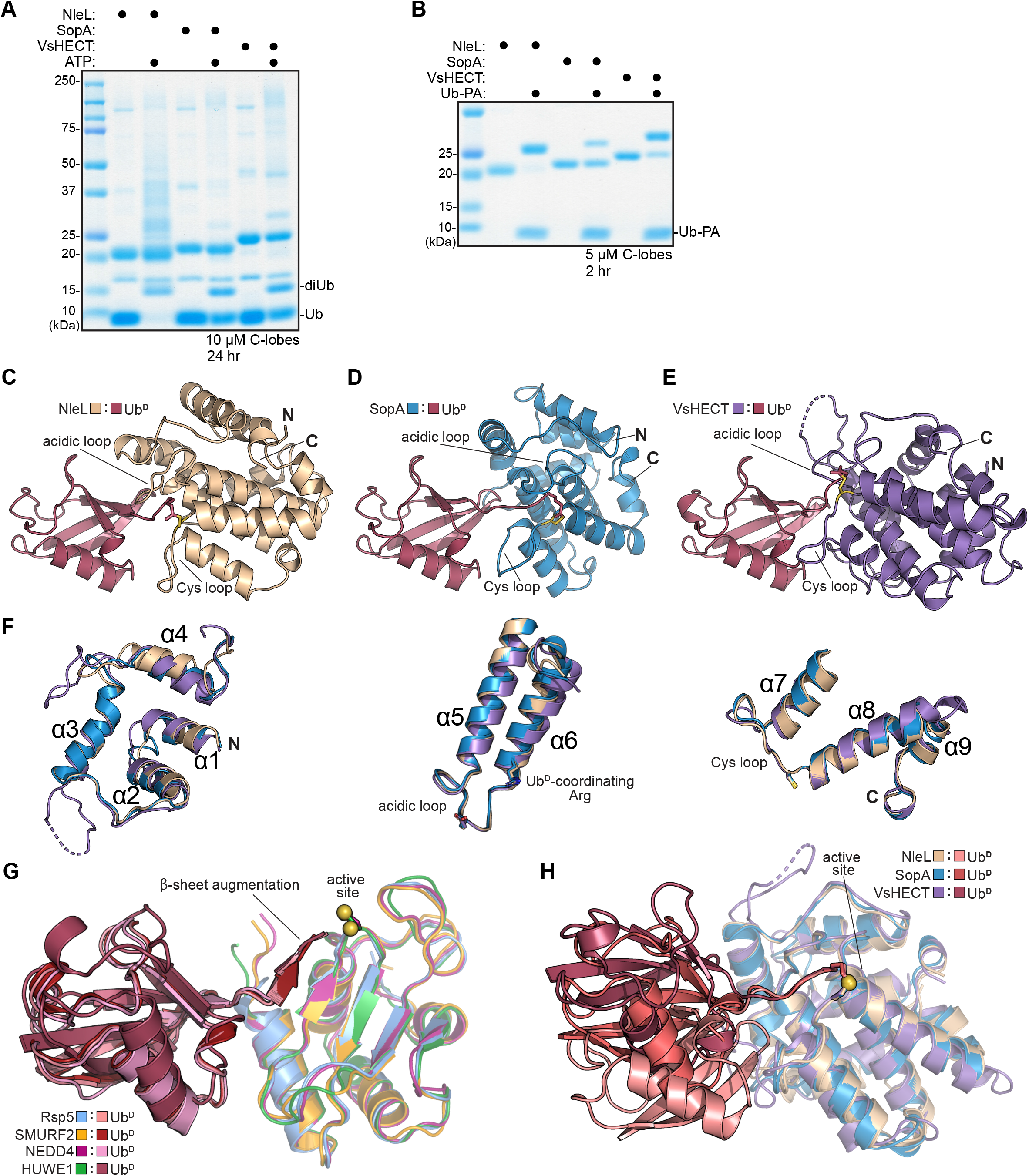
Structural and biochemical analysis of bHECT C-lobes. A. Gel-based Ub ligase assay of isolated bHECT C-lobe constructs. Reactions were initiated with ATP. bHECT concentrations are listed. Samples were quenched and resolved by SDS-PAGE with Coomassie staining. B. Gel-based reactivity assay using the Ub-PA probe with the isolated bHECT C-lobe constructs. bHECT concentrations are listed. Samples were taken at the indicated timepoints, quenched, and resolved by SDS-PAGE with Coomassie staining. C. 2.50 Å crystal structure of NleL-Ub^D^. The PA linkage at the active site Cys (yellow) is shown, and the N- and C-termini are labeled. Views in **C-E** were generated by aligning on Ub^D^. D. As in **C**, for the 1.75 Å SopA-Ub^D^ crystal structure. E. As in **C**, for the 1.44 Å VsHECT-Ub^D^ crystal structure. F. Overlay of the NleL, SopA, and VsHECT structures, aligned on the C-lobe and split into three sections to clearly show the conservation of each α-helical region. The α-helices are numbered starting from the N-terminus (labeled as “N”) to the C-terminus (labeled as “C”), with regions of interest (acidic loop, Cys loop, and critical residues) highlighted. G. Overlay of all available eHECT-Ub^D^ structures, aligned by the C-lobe portion of the HECT domain for NEDD4-Ub^D^ (PDB: 4BBN), HUWE1-Ub^D^ (PDB: 6XZ1), Rsp5-Ub^D^ (PDB: 4LCD), and SMURF2-Ub^D^ (PDB: 6FX4) with the active site Cys (yellow) highlighted. H. Overlay of the bHECT-Ub^D^ structures, aligned on their C-lobes, with the active site Cys (yellow) highlighted. See also Figure S2.

To obtain a better understanding of the bHECT Ub ligation pathway, we took advantage of the robust Ub-PA reactivity of the bHECT C-lobes and determined crystal structures for three complexes: NleL-Ub (2.50 Å), SopA-Ub (1.75 Å), and VsHECT-Ub (1.44 Å) (**Fig. 2C-E, S2A-C, Table 1**). Superposing the helical C-lobe domains of the bHECT-Ub^D^ structures revealed the overall similarity within each region of the fold (pairwise C-lobe Cα RMSD between 1.6 and 3.2 Å) (**Fig. 2F**). Although they adopt an α/μ structure distinct from bHECTs, eHECT C-lobes also demonstrate close structural homology to each other (pairwise Cα RMSD between 0.8 and 1.1 Å) (**Fig. 2G**). In contrast, while eHECT:Ub^D^ contacts are highly similar among resolved structures (pairwise Ub Cα RMSD between 0.7 and 5.7 Å)^18, 20, 22, 23^, the position of Ub^D^ on bHECT C-lobes is varied (pairwise Ub Cα RMSD between 8.0 and 15.4 Å) (**Fig. 2G-H**). When superposed onto previous apo NleL or SopA structures that encompass the μ-helix, N-lobe, and C-lobe domains, neither of the bound Ub^D^ molecules clash or form contacts with domains outside of the C-lobe (**Fig. S2D-E**).

**Table 1:** Data collection and refinement statistics.

|  | NleL-Ub | SopA-Ub | VsHECT-Ub |
| --- | --- | --- | --- |
| **Data collection** |  |  |  |
| Space group | I 1 2 1 | P 21 21 21 | P 1 21 1 |
| Cell dimensions |  |  |  |
| *a*, *b*, *c* (Å) | 76.269, 61.023, 116.188 | 51.893, 63.644, 81.409 | 35.855, 157.276, 53.025 |
| $\alpha$, $\beta$, $\gamma$ (°) | 90, 99.2508, 90 | 90, 90, 90 | 90, 93.756, 90 |
| Resolution (Å) | 38.5-2.50 (2.59-2.50)* | 36.06-1.75 (1.78-1.75) | 39.32-1.44 (1.46-1.44) |
| *R*<sub>merge</sub> | 0.131 (0.718) | 0.046 (0.665) | 0.036 (0.597) |
| *I* / $\sigma I$ | 6.2 (1.6) | 16.9 (2.00) | 17.7 (1.9) |
| Completeness (%) | 98.1 (96.8) | 98.7 (98.1) | 86.7 (41.7) |
| Redundancy | 3.1 (3.0) | 4.6 (4.5) | 3.9 (3.5) |
| **Refinement** |  |  |  |
| Resolution (Å) | 38.5-2.50 (2.59-2.5) | 36.06-1.75 (1.81-1.75) | 32.57-1.44 (1.49-1.44) |
| No. reflections | 37914 | 54714 | 91212 |
| *R*<sub>work</sub> / *R*<sub>free</sub> | 0.2036 / 0.2516 | 0.1765/0.1959 | 0.1699/0.1979 |
| No. atoms |  |  |  |
| Protein | 4055 | 2161 | 4649 |
| Ligand/ion | 8 | 4 | 22 |
| Water | 132 | 218 | 540 |
| *B*-factors |  |  |  |
| Protein | 40.11 | 28.67 | 22.83 |
| Ligand/ion | 36.03 | 29.45 | 16.92 |
| Water | 36.40 | 38.98 | 31.66 |
| R.m.s. deviations |  |  |  |
| Bond lengths (Å) | 0.009 | 0.007 | 0.012 |
| Bond angles (°) | 1.06 | 0.85 | 1.25 |

\*Values in parentheses are for highest-resolution shell.

### Donor Ub activation by bHECTs

Previous structural work for eHECTs NEDD4, HUWE1 and SMURF2 bound to Ub^D^ revealed a conserved coordination of the Ub^D^ C-terminal tail via several intersubunit contacts^18, 20, 22^. In the eHECT-Ub^D^ structures, residues 73-75 of the Ub^D^ C-terminus form a parallel β-strand with the conserved β-sheet of eHECT C-lobes, a feature referred to as β-sheet augmentation (**Fig. 2G, 3A**). Though they lack the β-sheet architecture, the bHECT C-lobes also exhibit a strong coordination of residues 73-75 from the Ub^D^ C-terminal tail, primarily through an extensive hydrogen bonding network (**Fig. S3A**). Coordination of the Ub^D^ C-terminal tail appears to primarily rely on a conserved bHECT Arg residue at the base of α-helix 6, which hydrogen bonds to the peptide backbone of Ub^D^ R74 (**Fig. 3B, S3B**). Mutation of this contact severely diminishes the ability of the bHECTs to ligate Ub in FP- or gel-based assays (**Fig. 3C-F**), and to react with the Ub-PA probe (**Fig. S3C**). NleL and SopA mediate secondary contacts to the Ub^D^ C-terminus via hydrogen bonds from E710 and D707, respectively, and mutations at these sites also reduce ligase activity (**Fig. 3C, E, S3A**). Thus, similar to eHECTs and other human ligase complexes^47–49^, bHECTs stretch and coordinate the C-terminal tail of Ub^D^, likely priming the C-terminus for nucleophilic attack by an incoming Lys.

**Figure 3:**
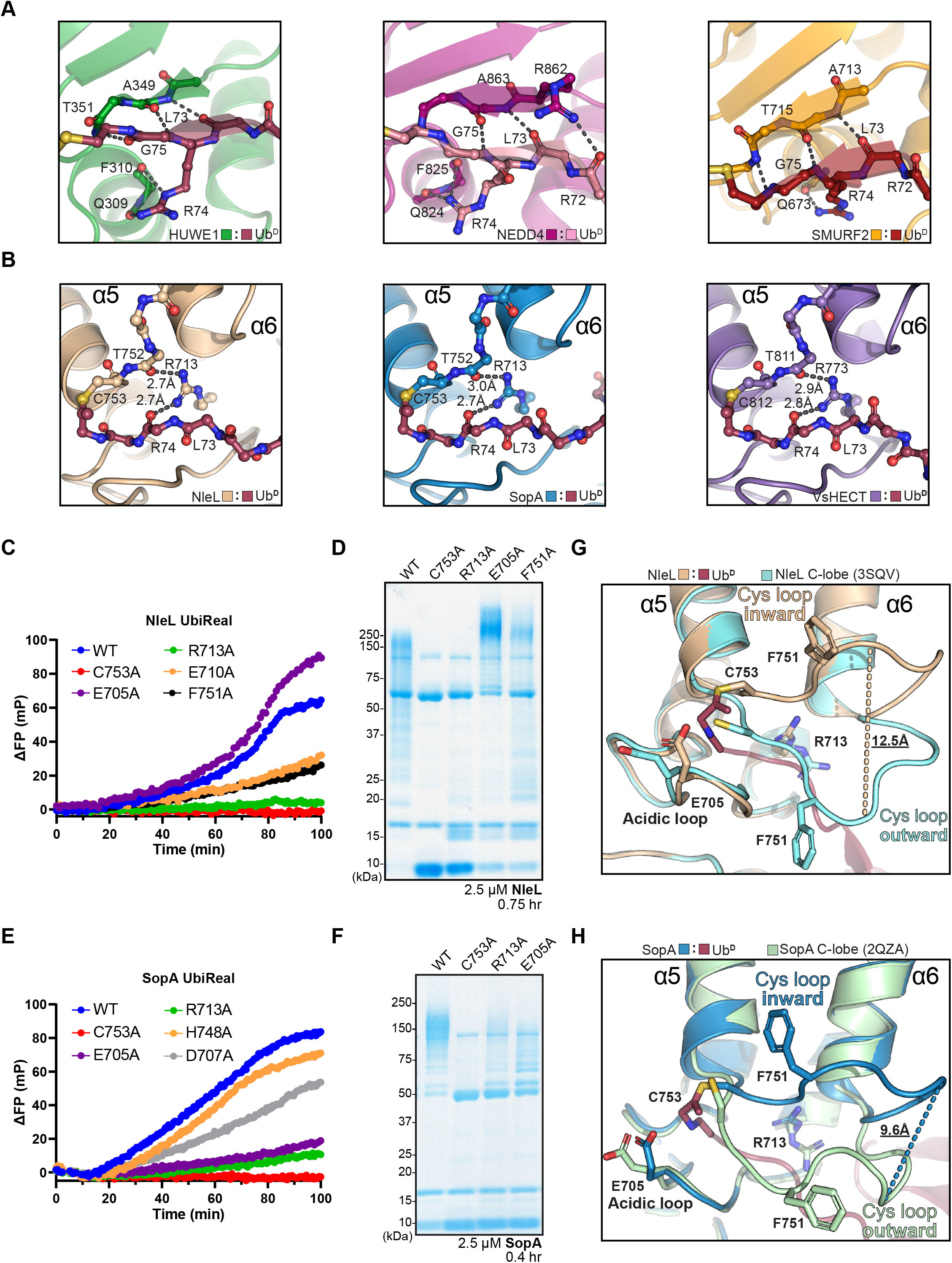
bHECT activation of Ub^D^. A. Beta-sheet augmentation between the Ub^D^ C-terminal tail and the eHECT C-lobes HUWE1-Ub^D^ (PDB: 6XZ1), NEDD4-Ub^D^ (PDB: 4BBN), and SMURF2-Ub^D^ (PDB: 6FX4). Hydrogen bonds between labeled residues are shown by black dashes. B. Ub^D^ C-terminal tail coordination by the conserved bHECT Arg residue at the base of α-helix 6 in NleL-Ub^D^, SopA-Ub^D^, and VsHECT-Ub^D^. Hydrogen bonds between labeled residues are shown by black dashes. C. Ubiquitin ligation assay monitored by the FP-based method UbiReal, for WT NleL and sequence- or structure-guided mutations at 2 µM. Reactions were initiated with ATP at timepoint 0 min. D. Gel-based Ub ligase assay of WT NleL and sequence- or structure-guided mutations. Reactions were initiated with ATP at timepoint 0 min. WT or mutant NleL were used at 2.5 µM and sampled at the indicated timepoints, quenched, and resolved by SDS-PAGE with Coomassie staining. E. As in **C**, for SopA constructs. F. As in **D**, for SopA constructs. G. Structural overlay highlighting the large movement of the Cys loop from the outward conformation observed in the apo NleL structure (PDB: 3NB2) to the inward conformation observed upon Ub^D^ binding to NleL. Some conserved residues of the Cys loop and acidic loop are shown. H. Structural overlay highlighting the large movement of the Cys loop from the outward conformation observed in the apo SopA structure (PDB: 2QYU) to inward conformation observed upon Ub^D^ binding to SopA. Some conserved residues of the Cys loop and acidic loop are shown. See also Figure S3.

Outside of contacts to the Ub^D^ C-terminal tail, we noted additional Ub^D^ contacts in the SopA and VsHECT structures. SopA forms multiple hydrogen bonds between H748 and E34 of Ub^D^, as well as a single hydrogen bond between H745 and T9 of Ub^D^ (**Fig. S3D**). A SopA H748A mutation showed a small effect on ligase activity by UbiReal (**Fig. 3E**). The VsHECT C-lobe featured unique contacts to both the I36 and L8 hydrophobic patches of Ub^D^, which were partly mediated by a unique insertion near the beginning of the C-lobe (**Fig. S3E-F**). Mutation of residues contacting either patch greatly reduced the ability of VsHECT to synthesize diUb (**Fig. S3G**). Altogether, while contacts at or near the Ub^D^ C-terminus are conserved and functionally required, additional contacts outside of the active site make important contributions to bHECT ligase activity as well.

Across all three bHECT-Ub^D^ structures, we noted that the Ub^D^ C-terminal tail was sandwiched between two loops: a “Cys loop” with a conserved Phe that precedes the active site Cys, and an “acidic loop”, which contains a conserved Glu residue that was previously proposed to play a catalytic role as a general base (**Fig. 2C-E, 3G-H, S3B**)^15^. Relative to the apo C-lobe structures, the Cys loops of both NleL and SopA undergo a substantial rearrangement upon linkage to Ub^D^ (**Fig. 3G-H**). The Cys loops of the apo bHECTs sit in an outward conformation, away from α-helices 5 and 6, while in all three Ub^D^-bound structures, the Cys loops tuck inward. This 12.5 Å and 9.6 Å rearrangement in NleL and SopA, respectively, coincide with rearrangements of the Ub^D^-coordinating Arg that position it to contact both the Ub^D^ C-terminus as well as the Cys loop backbone. The Glu residue of the acidic loop also adopts a conformation closer to the active site in the Ub-bound structures (**Fig. 3G-H**).

Considering the conformational changes upon Ub^D^ binding, we assessed the importance of impacted residues on bHECT ligase function. Within the NleL Cys loop, an F751A mutation greatly reduced ligase activity relative to WT (**Fig. 3C**). An NleL E705A mutation within the acidic loop actually gave a higher final FP value relative to WT (**Fig. 3C**). Using a gel-based readout, we observed that the NleL E705A mutant appeared to produce a higher molecular weight polyUb smear relative to NleL WT (**Fig. 3D**), which may partially explain the higher final FP value. Interestingly, in the case of SopA, the equivalent E705A mutation dramatically reduced activity (**Fig. 3E-F**). Thus, Cys loop and acidic loop residues appear to play important roles in bHECT ligase activity, but their precise functions were unclear from the bHECT-Ub^D^ structures alone.

### Model of E2-bHECT transthiolation

Previous work has determined crystal structures of NleL and SopA bound to the E2, UBE2L3^15^. We found that overlaying the NleL:UBE2L3 structure with our NleL-Ub^D^ structure yielded a feasible model for an E2:NleL∼Ub intermediate that occurs immediately following transthiolation of Ub to the E3, and before E2 dissociation (**Fig. 4A**). In this model, the orientation of E2 and Ub resemble a “backbent” conformation that has previously been observed among isolated E2∼Ub conjugates^50–54^. Within the E2:C-lobe interface in the published NleL:UBE2L3 structure, we noted a lack of electron density for UBE2L3 side chains in Loop 8, and a complete lack of electron density for the NleL Cys loop (**Fig. S4A**). In contrast, the NleL-Ub^D^ and NleL apo (as well as SopA-Ub^D^ and SopA apo) structures resolve the Cys loop in its inward and outward conformations (**Fig. 3G-H, S4B-C**), suggesting that the Cys loop is more dynamic in the NleL:E2 complex.

**Figure 4:**
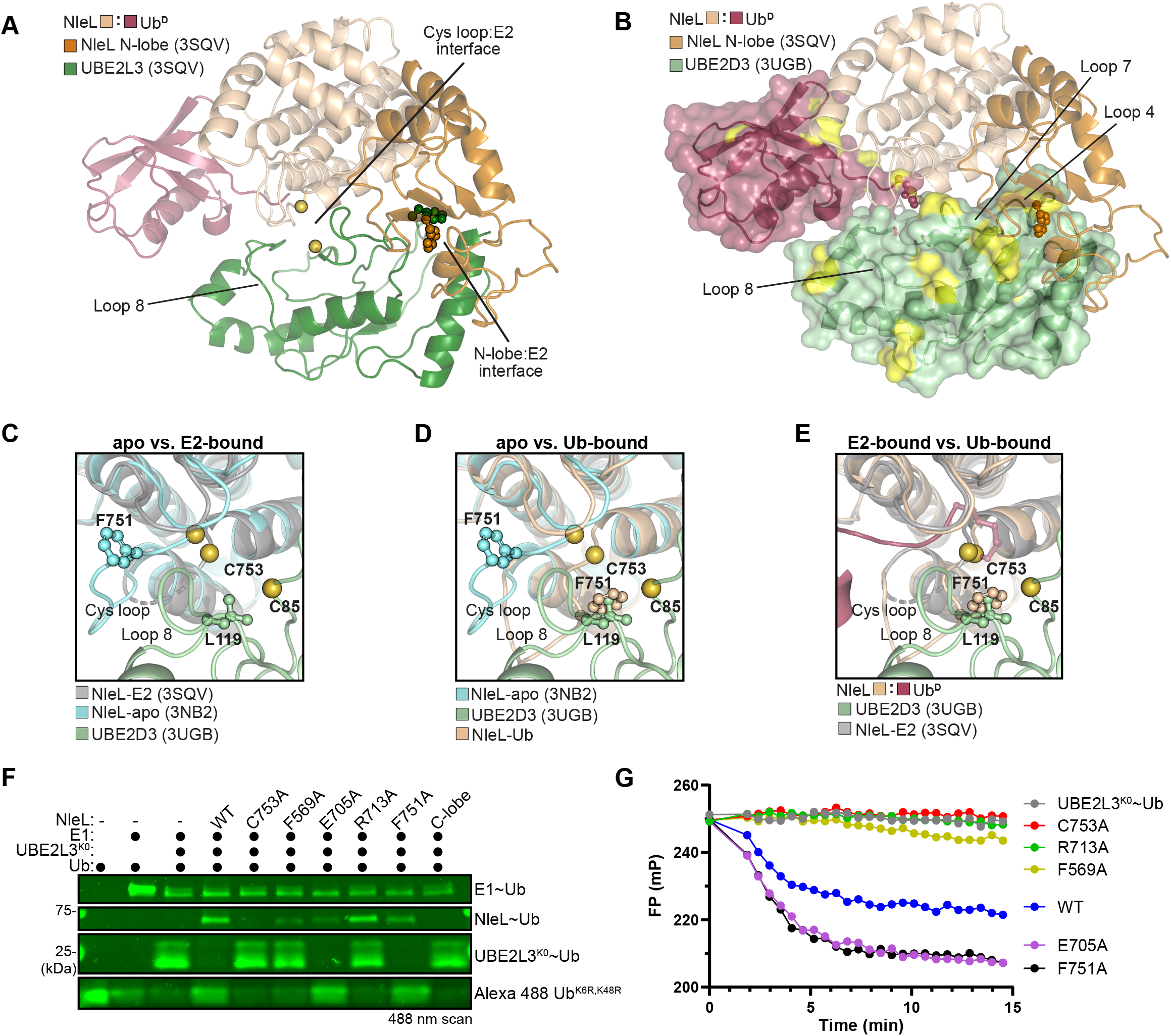
Model for E2-bHECT transthiolation. A. View of NleL-Ub^D^ and NleL:UBE2L3 (PDB: 3SQV) overlaid structures, representing a model of the E2:NleL∼Ub intermediate. View is obtained after aligning the two structures on the C-lobe of NleL, with only the C-lobe of the NleL-Ub^D^ structure shown. The E2:Cys loop and E2:N-lobe interfaces are highlighted. The conserved Phe residues at the E2:N-lobe interface are shown as sticks, and the active site Cys residues for both NleL and UBE2L3 and shown as yellow spheres. B. Structural model of the UBE2D3:NleL∼Ub complex with the significant peak intensity changes from **S4D-E** colored in yellow. The E2:N-lobe and active site interfaces are highlighted. C. View of NleL C-lobe Cys loops at the E2 interface comparing apo (PDB: 3NB2) and E2-bound (PDB: 3SQV) NleL structures. Note that the Cys loop could not be modeled in the E2-bound NleL structure and is shown in dashes. The Cys-loop Phe residue is shown for apo NleL. The active site Cys residues for NleL and UBE2L3 are shown as yellow spheres. Residue L119 of UBE2D3, near the C-lobe interface, is also shown. D. As in **C**, for the apo NleL (PDB: 3NB2) and NleL-Ub^D^ structures, highlighting the movement of the Cys loop from the outward conformation of the apo structure to the inward conformation of the NleL-Ub^D^ structure, and the resultant clash between the NleL-Ub^D^ Cys-loop Phe and residue L119 of UBE2D3 in the model. E. As in **C**, for the E2-bound (PDB: 3SQV) and NleL-Ub^D^ structures, highlighting the position of Ub^D^ at the interface of the E2:NleL∼Ub^D^ model. F. Gel-based transthiolation assay using Lys-less UBE2L3^K0^ and an N-terminally labeled Alexa 488 Ub K6,K48R substrate that prevents NleL from forming polyUb chains. EDTA was added after E2∼Ub formation to prevent recycling of the Ub. Slices of Ub, E1∼Ub, UBE2L3^K0^∼Ub, and NleL∼Ub (WT or mutant) from the same gel are shown for clarity. Samples were quenched in non-reducing sample buffer after reacting with the UBE2L3^K0^∼Ub for 5 min at 22 °C, resolved by SDS-PAGE, and scanned at 488 nm. G. E2∼Ub discharge assay monitored by the FP-based method UbiReal. N-terminally labeled Alexa 488 Ub K6,K48R substrate and Lys-less UBE2L3^K0^ were used to generate UBE2L3^K0^∼Ub conjugate prior to addition of buffer (control), NleL WT, or NleL mutants and subsequent measurement of FP changes. EDTA was added prior to the addition of NleL to prevent recycling of the Ub. See also Figure S4.

To verify our model of the NleL:E2:Ub interface in solution, we turned to NMR as a highly sensitive approach for studying transient protein interactions. We elected to study interactions with the well-characterized E2 UBE2D3, which is active with NleL and exhibits a high degree of structural homology to UBE2L3 (**Fig. S4A**)^14^. We generated a stable, monomeric UBE2D3-O-Ub conjugate by incorporating the UBE2D3 active site C85S mutation as well as the ‘backside’ S22R mutation. ^1^H,^15^N-TROSY spectra of ^15^N-labeled UBE2D3-O-Ub upon titration of either the NleL C-lobe alone or the full HECT-like domain revealed the interaction to be in the intermediate exchange regime resulting in selective peak broadening and intensity loss. Analysis of changes in peak intensities during the titration allowed identification of specific E2∼Ub residues involved in binding to NleL (**Fig. S4D-E**). Resonances that exhibited a significant reduction in peak intensity were mapped onto a surface representation of UBE2D3 and Ub within the modeled complex (**Fig. 4B**). The results were consistent with interactions to the N- and C-lobes of NleL in our model. The resonance corresponding to F62, the UBE2D3 residue in Loop 4 critical for interaction with the NleL N-lobe, broadened significantly with titration of both the full NleL HECT-like domain and the isolated C-lobe construct (**Fig. 4B, S4D-E**). In our model, the NleL C-lobe does approach UBE2D3 underneath Loop 4, and the aromatic nature of F62 might make it particularly sensitive to reporting on this interaction. In contrast, significant peak broadening was observed for Loop 7 (residues 90-95) of UBE2D3 only in the presence of the N-lobe, which can be explained in our model by contacts from an NleL loop downstream of the conserved F569. Significant peak broadening within the Ub C-terminal tail was also observed with titration of the full HECT-like domain, consistent with contacts to the NleL C-lobe prior to transthiolation (**Fig. 4B, S4D**).

Using our validated model for Ub transthiolation, we sought to interpret how conformational changes in the NleL Cys loop may impact E2 binding. In the apo NleL structure, the Cys loop sits in the outward orientation, away from the E2 interface (**Fig. 3G, 4C**). Upon binding of Ub^D^ and the subsequent rearrangement of the Cys loop to the inward conformation, the Cys loop, and in particular F751, clashes with Loop 8 the E2 (**Fig. 4D**). However, the Ub^D^ itself doesn’t appear to clash at the NleL Cys loop:E2 interface (**Fig. 4E**). Altogether, this suggests the Cys loop rearrangement to the well-ordered inward conformation following Ub transthiolation may result in steric clashes that help to dissociate the C-lobe from the E2, though not necessarily breaking E2:E3 interactions within the N-lobe. This would be consistent with the two different C-lobe conformations that are observed between the apo and E2-bound NleL structures^14, 15^.

Since important residues in the Cys loop and acidic loop are located near the modeled E2:C-lobe interface, we sought to test whether their mutation impacted transthiolation from the E2 (e.g., discharging the E2∼Ub bond to form E3∼Ub or free Ub). We first generated an E2∼Ub conjugate between Lys-less UBE2L3^K0^ (to prevent E2 ubiquitination), and a fluorescently-labeled Ub that contained K6R and K48R mutations (to prevent polyUb chain formation). NleL WT completely discharged the E2∼Ub conjugate to generate E3∼Ub or free Ub, while the catalytically inactive NleL C753A failed to do so (**Fig 4F**). This indicated that E2∼Ub discharge was dependent on transthiolation to the NleL active site Cys, and that any released Ub from the reaction was a result of discharge from the E3∼Ub intermediate. Consistent with this model, an F569A mutation within the N-lobe E2-binding site showed very minor discharge of E2∼Ub and formation of E3∼Ub, while an isolated C-lobe construct showed no E2∼Ub discharge. Consistent with a role in activating the Ub^D^ C-terminus (**Fig. 3B**), the NleL R713A mutant could still receive Ub from the E2 but was very inefficient at discharging it. For both the Cys loop mutant F751A and the acidic loop mutant E705A, complete discharge of the E2∼Ub conjugate was observed, primarily yielding free Ub. In contrast to NleL WT, the E3∼Ub intermediate was not observed (**Fig. 4F**). This indicated that transthiolation from the E2∼Ub did not appear to be inhibited, and the resulting E3∼Ub conjugate formed by these mutants may be more labile toward hydrolysis than WT. A modified FP-based UbiReal assay was used to corroborate these observations with better temporal resolution (**Fig. S4F**). Monitoring fluorescent Ub incorporated into an E2∼Ub conjugate, the E705A and F751A mutants produced lower FP values, matching results from the gel-based assays indicating a larger ratio of free Ub to E3∼Ub intermediate as compared to NleL WT (**Fig. 4G**). Furthermore, the steady FP signals of the NleL C753A and NleL R713A reactions indicated an inability of these mutants to discharge the E2∼Ub conjugate.

### NleL coordination of K48 acceptor Ub

HECTs, as well as other Cys-based Ub ligases, have the capability to preferentially generate one or several different types of polyUb linkages. How HECT domains coordinate an acceptor Ub for linkage-specific ligation is largely enigmatic. During our analysis of the NleL-Ub^D^ structure, we observed close crystal contacts between NleL-Ub^D^ active sites and Ub K48 from neighboring molecules representing a potential acceptor ubiquitin, Ub^A^ (**Fig 5A-B, S5A**). NleL is known to catalyze a mixture of K6- and K48-linked polyUb^14, 34, 55^, but as a first step toward interpreting the NleL:Ub^A^ interface we tested if polyUb specificity is retained within the C-lobe construct that was crystallized. Using a K-only panel of Ub mutants, in which all Lys residues but one had been mutated to Arg, we observed that the NleL C-lobe construct preferentially generated K6- and K48-linked polyUb, in accordance with previous data for the full HECT domain^14, 34^ (**Fig. S5B**). The specificity of the SopA C-lobe construct toward K48-linked polyUb was also consistent with previous data^17^ (**Fig. S5C**), indicating that bHECT C-lobes represent a minimal unit for polyUb linkage specificity. Thus, in our structure of the NleL-Ub^D^ intermediate, we fortuitously captured a snapshot of K48 polyUb ligation.

**Figure 5:**
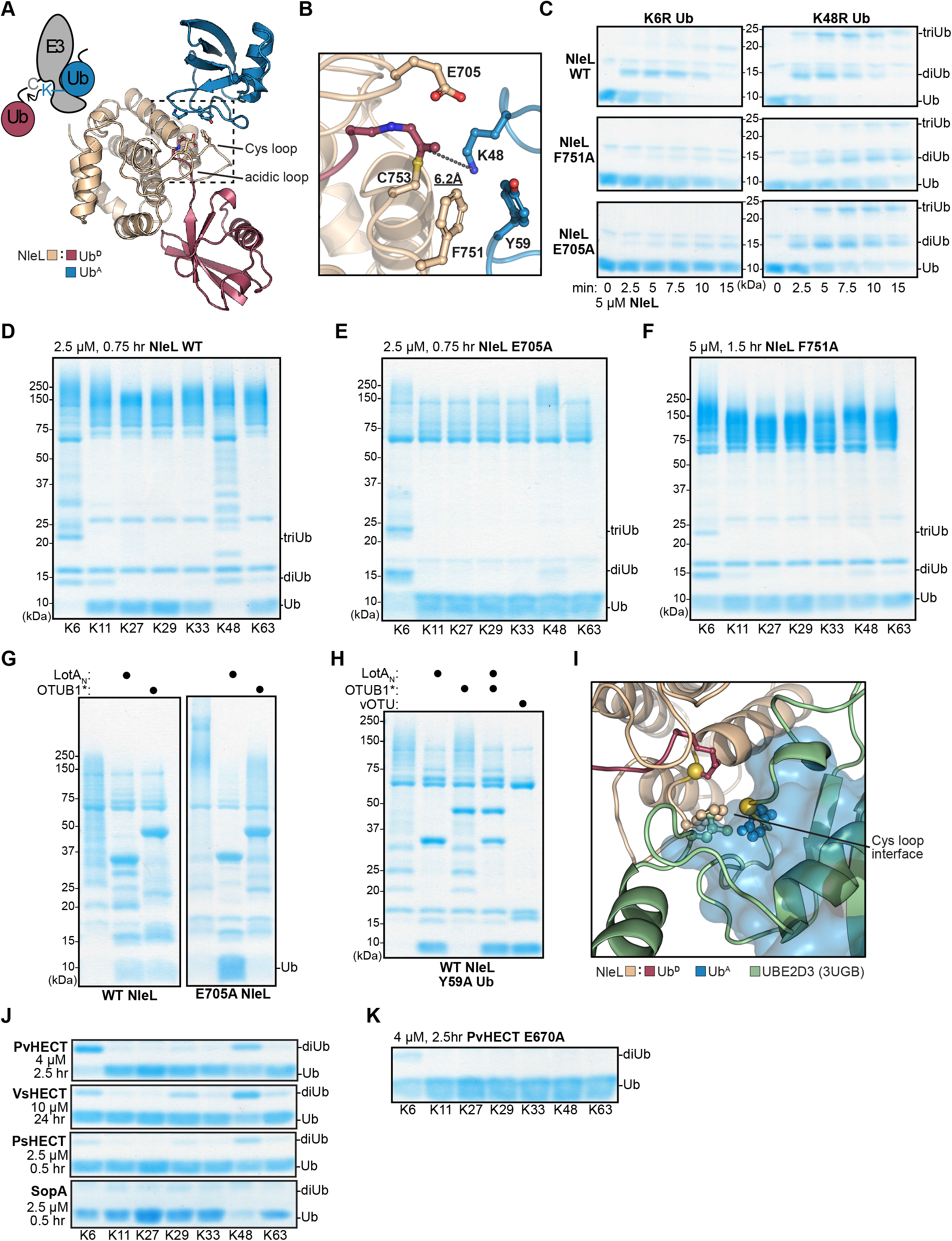
bHECT coordination of an acceptor Ub. A. View of the Ub^A^:NleL-Ub^D^ interface observed through crystal symmetry, with key residues at the interface highlighted. A cartoon depiction of diUb ligation by a HECT ligase is shown for comparison. B. Zoomed-in view of the Ub^A^:NleL-Ub^D^ interface shown in **A**, with key residues highlighted. The distance between the ε-amino group of K48 and the Ub^D^ C-terminus is shown. C. Gel-based assay monitoring the consumption of K6R or K48R Ub by NleL WT, NleL F751A, and NleL E705A. Reactions were sampled at the indicated timepoints, quenched, and resolved by SDS-PAGE with Coomassie staining. D. Gel-based polyUb specificity assay for NleL WT using the panel of K-only Ub mutants, each containing only the single Lys indicated with all others mutated to Arg. Reactions were quenched and resolved by SDS-PAGE with Coomassie staining. E. As in **D**, for the NleL E705A mutant. F. As in **D**, for the NleL F751A mutant. G. UbiCRest assay monitoring the cleavage of polyUb generated by NleL WT or NleL E705A using K6-specific LotA_N_ and K48-specific OTUB1*. DUB-treated and control samples were quenched and resolved by SDS-PAGE with Coomassie staining. H. As in **G**, for polyUb generated by NleL WT with Ub Y59A. In addition to LotA_N_ and OTUB1*, the nonspecific DUB vOTU is used for comparison. I. Structural overlay showing overlap of the Ub^A^- and UBE2D3-binding sites on the NleL C-lobe. Important interface residues are shown. J. As in **D** for PvHECT, VsHECT, PsHECT, and SopA. Only the monoUb and diUb region of the gels are shown for clarity. K. As in **D** for the PvHECT E670A acidic loop mutant. See also Figure S5.

In addition to Ub^A^ K48 approaching the NleL active site C753, we observed several other notable contacts at the NleL:Ub^A^ interface. Residue F751 of the NleL Cys loop, positioned in the inward conformation following conjugation of Ub^D^ to the NleL active site (**Fig. 3G**), forms a hydrophobic interface with Y59 of the Ub^A^ (**Fig. 5B**). As for the acidic loop, residue E705 that was observed to approach the active site upon Ub^D^ conjugation (**Fig. 3G**), is also near the NleL:Ub^A^ interface (**Fig. 5B**). Since only the F751A mutant affected total ligase activity and neither mutant affected E2-NleL transthiolation (**Fig. 3C-F, 4F-G**), we tested if these residues were involved in K48-specific polyUb ligation by NleL. We first established comparative NleL ligation reactions using K6R or K48R Ub as substrates, producing K48- and K6-linked polyUb, respectively (**Fig. 5C**). While NleL WT consumed the Ub substrates at equal rates, both the F751A and E705A mutants greatly preferred the K48R substrate, and were very slow to produce any polyUb products with the K6R substrate (**Fig. 5C**). The F751A mutant was markedly slower than WT to produce polyUb with the K48R substrate, suggesting that this region of the NleL Cys loop may also play a role in assembly of K6 polyUb. Next, we analyzed polyUb specificity of the NleL mutants using the panel of K-only Ub mutants. Remarkably, the E705A mutation severely abrogated the ability of NleL to generate K48 polyUb relative to WT, rendering it largely specific for K6 polyUb (**Fig. 5D-E**). The F751A mutation also inhibited K48 polyUb ligation in this assay, though total Ub ligation also appeared to be impaired (**Fig. 5F**).

PolyUb specificity with a native Ub substrate was validated using UbiCRest, an assay that uses linkage-specific deubiquitinating enzymes (DUBs) to determine the types of polyUb linkages present in a sample^56^. To distinguish between K6- and K48-linked polyUb, we utilized the recently described K6-specific DUB LotA^N^ from *Legionella pneumophila*^57, 58^, as well as the optimized human K48-specific DUB, OTUB1*^59^. PolyUb chains generated by WT NleL were cleaved equally well by both LotA_N_ and OTUB1*, yielding similar amounts of released monoUb (**Fig. 5G**). However, polyUb chains generated by NleL E705A were more robustly cleaved by LotA_N_, which is especially apparent when looking at the return of free monoUb (**Fig. 5G**). The role of E705 in K48-linked polyUb ligation is also consistent with the SopA E705A mutant, which shows a more substantial defect in total ubiquitination, likely because it favors just the single polyUb linkage type (**Fig. 3E-F, S5C**). Testing the opposite side of the NleL:Ub^A^ interface, incorporation of a Ub Y59A mutation ablated the ability of WT NleL to produce K48-linked polyUb without affecting assembly of K6-linked polyUb (**Fig. 5H**). Interestingly, Y59 of Ub^A^ occupies a similar position as UBE2D3 L119 in the modeled UBE2D3:NleL∼Ub complex, suggesting that the E2 must either dissociate from the N-lobe, or the C-lobe must rearrange to a new conformation in order to allow K48-linked polyUb ligation (**Fig. 5I**).

Since the polyUb specificity of NleL can be redirected with single point mutations, we examined if these features directing linkage specificity were shared by other bHECTs (**Fig. 1C**). We monitored disappearance of the K-only Ub substrates and formation of diUb to profile polyUb specificity. Across the panel of bacterial HECT-like ligases, there was an underlying trend to ligate K6- and K48-linked polyUb to varying extents (**Fig. 5J**). SopA preferentially generated K48-linked polyUb, as previously established. VsHECT and PsHECT appeared to prefer K48-linked polyUb ligation, though some other linkages were observed as well. Interestingly, PvHECT appeared to natively prefer K6 ligation, despite a conserved Glu on the acidic loop and a Phe on the Cys loop (**Fig. 5J, S3B**). This could indicate that the putative K6 Ub^A^ acceptor site of PvHECT may have a higher binding affinity than its K48 Ub^A^ acceptor site. Mutating the PvHECT acidic loop Glu residue, analogous to NleL E705, also inhibited formation of the residual K48 linkages, though overall ligase activity appeared to be impaired as well (**Fig. 5J-K**).

Previous work has shown that some eHECT C-lobes can be swapped to alter polyUb specificity^18, 30^. Due to the conserved fold among bHECT C-lobes (**Fig. 2F**), and because bHECT polyUb specificity is fully encoded within the C-lobe (**Fig. S5B-C**), we hypothesized that replacing the C-lobe of SopA with that of NleL would rewire SopA’s ligase activity (**Fig. S5D**). Using the K-only panel of Ub mutants, we observed that the SopA-NleL chimera (SNc) ligase was able to ligate both K6- and K48-linked polyUb, similar to NleL (**Fig. S5E**). Further, adding the E705A acidic loop mutation eliminated most K48 ligation, resulting in a SopA construct rewired for K6-linked polyUb (**Fig. S5E**).

### HUWE1 polyUb specificity augmentation

The structural and biochemical work reported above illustrate clear roles for Cys loop and, in particular, acidic loop residues in controlling bHECT polyUb specificity. Though topologically different, this dual loop architecture is also present in eHECTs, wherein the active site Cys sits together with a Phe on a loop connecting two β-strands and is positioned adjacent to an acidic loop containing a Glu/Asp residue (**Fig. 6A-B, S6A**). Although the context may not be conserved, we hypothesized that a cryptic acidic loop may still be important for eHECT polyUb specificity. Aligning the Ub^D^ C-terminal tails across the HUWE1-Ub^D^ and the Ub^A^:NleL-Ub^D^ structures placed the Ub^A^ in a plausible orientation for HUWE1-Ub^D^ ligation and highlighted the proximity to the putative acidic loop residue E4315 (**Fig. S6B**). As previous work indicated a reliance on the N-lobe for Ub recognition^23^, we additionally expanded the search beyond the C-lobe for acidic loops, utilizing previously determined structures of the apo or Ub-bound HUWE1 HECT domain^23, 25^. Analysis of these structures revealed two additional potential acidic loops (**Fig. 6C, S6C**). In the HUWE1-Ub^D^ structure, which captures the “L” conformation of the HECT domain, an acidic loop from the N-lobe encoding E4054 and Q4056 is in close proximity to the active site (**Fig. 6C**). Interestingly, this loop matches by sequence and structural alignment to a structurally unresolved loop of Rsp5 that was previously demonstrated to have a critical catalytic function (**Fig. S6D**)^22^. In the apo HUWE1 structure, the C-lobe is shifted into a “T” conformation that positions a different N-lobe acidic residue, D4087, near the active site (**Fig. S6C**).

**Figure 6:**
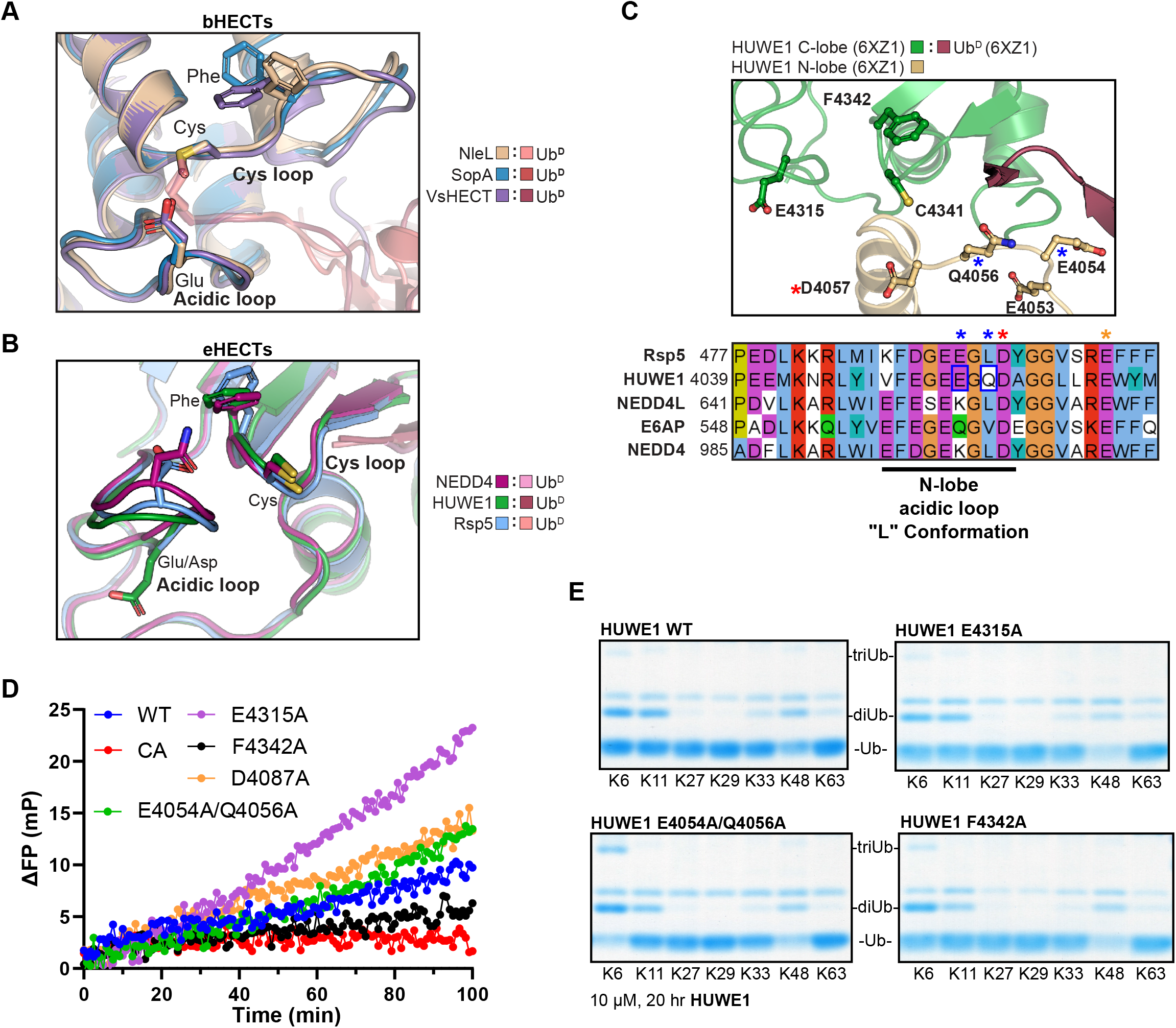
HUWE1 mutants show increased K6 Ub ligation. A. Overlay of the bHECT C-lobes, emphasizing the orientation of the Cys loop and acidic loop at the bHECT:Ub^D^ interface for NleL, SopA, and VsHECT. Residues that are structurally conserved between bHECTs and eHECTs are shown. B. Overlay of the eHECT C-lobes, emphasizing the orientation of the Cys loop and acidic loop at the eHECT:Ub^D^ interface for NEDD4, HUWE1, and Rsp5. Residues that are structurally conserved between bHECTs and eHECTs are shown. C. Structure of eHECT HUWE1-Ub^D^ (PDB: 6XZ1), focusing on the active site, with the C-lobe shown in green and the N-lobe shown in gold. The C-lobe acidic loop containing E4315 is shown, as well as an additional acidic loop from the L conformation of the N-lobe. Sequence conservation of the N-lobe acidic loop is shown with other eHECTs. The location of an Rsp5 acidic residue previously shown to be important for activity is indicated by a red star. The location of the eHECT E6AP Glu residue (not shown in the structure panel) mutated in Angelman’s syndrome is indicated by an orange star. HUWE1 sites selected for mutational analysis are indicated with blue boxes and blue stars. D. E3 ligase assay monitored by the FP-based method UbiReal, for WT HUWE1 and the sequence- or structure-guided mutants at 25 µM. Reactions were initiated with ATP at time point 0 min. E. Gel-based polyUb specificity assay for HUWE1 WT, the N-lobe acidic loop mutant E4054A/Q4056A, the C-lobe acidic loop mutant E4315A, and the Cys loop mutant F4342A, using the panel of K-only Ub mutants. Reactions were quenched and resolved by SDS-PAGE with Coomassie staining. Gel regions corresponding to monoUb, diUb, and triUb are shown for clarity. See also Figure S6.

We used the HUWE1 structures to guide mutations in the putative acidic loops, including HUWE1 E4315A, HUWE1 E4054A/Q4056A, and HUWE1 D4087A, and tested their effects on total activity in a UbiReal ligase assay. We also tested the Cys-loop Phe residue of HUWE1, F4342, as the structurally analogous residue of NleL contributed to polyUb specificity (**Fig. 5F**). Except for the C-lobe acidic loop mutation, E4315A, which appeared to increase activity, none of the acidic mutants appreciably altered ligase activity by this assay (**Fig. 6D**). Similar to what was observed in bHECTs, the HUWE1 Cys-loop mutant F4342A showed reduced overall ligase activity (**Fig. 6D**). Next, we assessed the mutational effects on polyUb specificity using the panel of K-only Ub mutants. Neither the C-lobe acidic mutant, E4315A, nor the T conformation acidic mutant, D4087A, had appreciable effects on polyUb specificity (**Fig. 6E, S6E**). The Cys-loop F4342A mutant had a minor impact on specificity, producing less K11-linked polyUb (**Fig. 6E**). Strikingly, however, the L conformation acidic mutant, E4054A/Q4056A, produced considerably more K6-linked polyUb and nearly consumed the available Ub substrate (**Fig. 6E, S6E**). Thus, residues within the eHECT N-lobe can contribute to ligase specificity, raising the possibility that distinct conformations of the HECT domain can influence the nature of the polyUb produced.

## DISCUSSION

Together with prior studies, our structural and biochemical data provide a complete picture of the bHECT ubiquitination reaction. Combining our Ub-activated NleL structure with a previous E2-bound structure yielded a composite model for the initial E2-E3 transthiolation reaction that is supported by NMR and biochemical data. Held in place by contacts to the N-lobe, the E2∼Ub conjugate is engaged by the bHECT C-lobe from the same direction as eHECTs, but opposite to eukaryotic RBR and RCR E3 ligases^15, 21, 60, 61^. Ub transfer onto the E3 active site is coincident with a large conformational rearrangement of the Cys loop, including a conserved Phe residue, that may act in part to displace the activated C-lobe. Among the bHECT-Ub structures that we determined, contacts made to the Ub β-grasp domain are highly variable, resulting in large differences in how the activated Ub is oriented. In contrast, the Ub C-terminus is stabilized in an extended conformation by a conserved group of hydrogen bonds, many of which arise from a bHECT Arg residue that is required for priming the donor Ub. Flexibility within the linker domain allows movement of the activated C-lobe toward the substrate for Ub transfer. Alternatively, bHECTs can assemble linkage-specific polyUb chains through an acceptor Ub-binding site, which is captured in one of our structures through crystal packing. The same Cys loop rearrangement that displaced the E2 also creates a Ub^A^-binding site, wherein the conserved Phe contacts Y59 of the incoming Ub, orienting its K48 toward the active site. This interface is essential, as mutating either side severely affects the ability of NleL to ligate K48-linked polyUb chains, with minimal or no effect on activity toward K6-linked polyUb. This structure provides the first glimpse of K48-specific polyUb ligation in any system and, interestingly, reliance upon Ub Y59 may be a common strategy for specificity, as the E2 enzymes UBE2K and UBE2R1 also require this contact^62, 63^. Across the NleL active site lies a conserved acidic loop, the mutation of which also toggles NleL activity away from K48 and toward K6-linked polyUb.

Through expansion of the bHECT family, we gained a better appreciation of its sequence and functional diversity. Remarkably, NleL is not alone in its ability to ligate atypical K6-linked polyUb, in fact it appears to be the preferred product of PvHECT from the opportunistic pathogen *P. vulgaris*. *Proteus* species are commonly associated with urinary tract infections, where they can form large extracellular clusters^64, 65^. EHEC also maintains an extracellular niche, the regulation of which has been tied to NleL ligase activity^16, 35^, suggesting that perhaps ligation of K6-linked polyUb plays a role for extracellular bacteria that is not required for the intracellular *Salmonella* Typhimurium, which encodes the K48-specific SopA. This raises an interesting contrast to recent work on other intracellular bacteria, such as *Legionella pneumophila*, which secrete DUBs that specifically remove K6-linked polyUb signals^57, 58, 66–69^. The signaling roles for K6-linked polyUb remain very murky, particularly with respect to the host-pathogen interface. Our newfound ability to modulate the polyUb specificities of bHECTs will provide important tools for future studies on this mysterious signal.

Despite their apparent differences in sequence and structure, many of the lessons learned from studying bHECTs could be translated to eHECTs. In particular, both bHECTs and eHECTs coordinate an extended C-terminal tail of Ub^D^, which is accomplished by β-sheet augmentation in the eHECTs^18, 20, 23^, and primarily through a conserved Arg in the bHECTs. Though the importance of these backbone interactions is difficult to test in eHECTs, we could show in bHECTs that mutation of the conserved Arg residue severely reduces ligase activity, presumably through an inability to orient the donor Ub for nucleophilic attack. We also observed that the Ub^D^ C-terminal tail is sandwiched between a Phe-containing Cys loop and an acidic loop for both eHECTs and bHECTs. Our structural work captured the importance of these loops in establishing an acceptor Ub-binding site, and while defining the basis of polyUb specificity among eHECTs has been a longstanding challenge, we could show that analogous loops in human HUWE1 also regulate polyUb specificity. Surprisingly, the HUWE1 acidic loop that influenced polyUb specificity to the largest extent was not encoded near the active site in the C-lobe, but was contributed from the N-lobe. This loop, by sequence and structure, corresponds to the location of an Asp residue critical for Rsp5 ligase activity^22^. Thus, for both SopA and Rsp5, which specifically ligate a single type of polyUb, mutation of the acidic loop ablates activity whereas for NleL and HUWE1, both of which encode multiple polyUb specificities, it instead alters the preferred product. This suggests the possibility that distinct acidic residues enable the formation of different polyUb products. In fact, many eHECTs encode conserved acidic residues near their C-termini, which are already known to partly mediate polyUb specificity in several _cases18,23,30,37._

The roles of acidic residues in Ub transfer are well documented, with mutations in the catalytic base generally resulting in deficient polyUb synthesis and mutations in the catalytic acid resulting in more stable E3∼Ub intermediates^22, 48, 49, 70, 71^. In general, acidic residues near the active site may function to deprotonate the ε-amino group of an incoming Lys on the acceptor Ub or a substrate, or simply guide the target Lys into the E3 active site. Remarkably, this underlying principle of Ub ligation is even followed by the most structurally distinct bacterial E3 ligases, including the Novel E3 Ligase (NEL) family found in *Salmonella* and *Shigella* species, as well as the SidC E3 ligase family from *Legionella* species^49, 70, 72, 73^. In the NEL family, mutation of a conserved Asp near the active site Cys resulted in retained E3∼Ub formation but deficient polyUb synthesis^74^. A second family member was shown to rely on two separate Asp residues, one acting as a catalytic base to deprotonate the incoming Lys and the second as a catalytic acid to support the tetrahedral intermediate^70^. SidC was also observed to encode two conserved Asp residues near the active site, both of which contribute to polyUb synthesis^73^. Clearly, despite large differences in structure and evolutionary convergence of Ub ligase function, certain principles of Ub transfer still hold true. Just as our work on bHECT E3 ligases has demonstrated for polyUb specificity, studying the principles of bacterial E3 ligases may yet reveal further insights into the mechanisms governing eukaryotic Ub biology.

## METHODS

### Bacterial HECT-like domain prediction

T-coffee^75^ was used to generate a consensus sequence from a multiple sequence alignment of the only two known HECT-like domains, NleL and SopA. With the consensus sequence of either the C-lobe alone, or the consensus sequence of the full HECT domain, the NCBI protein BLAST suite was used to search bacterial genomes for similar sequences. Sequences of bacterial proteins with HECT-like similarities were manually curated from BLAST by inspection for alignment to critical HECT-like features of NleL and SopA. Sequence features included an active site Cys residue, an E2-interacting aromatic residue, a linker region, and a HECT-like domain of similar size to NleL and SopA (∼400 residues. Candidate sequences were next subjected to protein homology modeling using Phyre2^76^. Protein models of the candidate sequences were aligned with structures of NleL and SopA in PyMol, and manually inspected for the bi-lobal structures characteristic of HECT and HECT-like domains. Candidates that met these criteria were synthesized (IDT), using codons optimized for *Escherichia coli* expression systems.

### Cloning and mutagenesis

The *nleL* gene was cloned from *Escherichia coli* O157:H7 str. Sakai, the *sopA* gene was cloned from *Salmonella enterica* Typhimurium SL1344, and all other bHECT constructs (VsHECT, PvHECT, PsHECT, and EaHECT) were synthesized by IDT (**Table S1**). All bHECT expression constructs were designed using Phyre2^76^ and the available crystal structures of NleL^14^ and SopA^13^. HUWE1 and E6AP were a kind gift from Thomas Mund (MRC Laboratory of Molecular Biology). All HECTs were cloned into the pOPIN-B vector which contains an 3C-cleavable N-terminal His-tag, except for EaHECT and E6AP, which were cloned into the pOPIN-S vector which additionally has an N-terminal SUMO tag. Cloning and mutagenesis were performed using Phusion DNA Polymerase (New England BioLabs) and TOP10 *Escherichia coli* (MilliporeSigma).

### Protein expression and purification

All pOPIN-B/S bHECT and eHECT constructs were expressed and purified similarly. Transformed Rosetta (DE3) *Escherichia coli* were grown in Luria broth containing 35 µg/mL chloramphenicol and 50 µg/mL kanamycin at 37 °C until OD_600_ 0.6-0.8, induced with 300 µM IPTG, and left to express at 18 °C for 18-20 hours. Cells were harvested by centrifugation and resuspended in 25 mM Tris, 200 mM NaCl, 2 mM β-mercaptoethanol, pH 8.0 (Buffer A). Following a freeze-thaw cycle, cells were incubated for 30 min on ice with lysozyme, DNase, PMSF, and SigmaFAST protease inhibitor cocktail (MilliporeSigma), then lysed by sonication. Clarified lysates were applied to HisPur cobalt affinity resin (ThermoFisher), washed with Buffer A containing 500 mM NaCl and 5 mM imidazole, and eluted using Buffer A containing 300 mM imidazole. bHECT and eHECT proteins were concentrated using Amicon centrifugal filters (MilliporeSigma) and applied to a HiLoad Superdex 75 pg 16/600 size exclusion column (Cytiva) equilibrated in 25 mM Tris, 150 mM NaCl, 0.5 mM DTT, pH 8.0 at 4 °C. Fractions were evaluated for purity by SDS-PAGE, collected, concentrated, and quantified by absorbance (280 nm) prior to flash freezing and storage at -80 °C.

Untagged WT or mutant Ub constructs were expressed from the pET-17b vector. Transformed Rosetta (DE3) *Escherichia coli* were grown by auto-induction in a modified ZYM-5052 media^77^ containing 35 µg/mL chloramphenicol and 100 µg/mL ampicillin at 37 °C for 24-48 h. Cells were harvested by centrifugation, resuspended, and lysed as above. Clarified lysates were acidified by dropwise addition of 70% perchloric acid to a final concentration of 0.5%. The mixture was stirred on ice for 1 h prior to centrifugation. The clarified supernatant was dialyzed into 50 mM sodium acetate, pH 5.0 overnight. The protein was applied to a HiPrep SP FF 16/10 cation exchange column (Cytiva), washed with additional 50 mM sodium acetate, pH 5, and eluted over a linear gradient to a matched buffer containing 500 mM NaCl. Ub was finally purified by application to a HiLoad Superdex 75 pg 16/600 size exclusion column equilibrated in 25 mM Tris, 200 mM NaCl, pH 8.0. Purified Ub was quantified by absorbance (280 nm), or by a BCA standard curve for Ub Y59A (ThermoFisher), and flash frozen for storage at either -20 °C or -80 °C.

^15^N-labeled proteins were grown in minimal MOPS medium supplemented with ^15^NH_4_Cl. ^15^N-Ub was expressed and purified as above for unlabeled Ub. Untagged ^15^N-UBE2D3 C85S/S22R was expressed from pET17b using IPTG induction as described above, harvested, and resuspended in 50 mM MES, pH 6.0. Cells were lysed by sonication as described above, and UBE2D3 was purified by cation exchange chromatography on a HiPrep SP FF 16/10 column (Cytiva) using a 0-500 mM salt gradient in 50 mM MES, pH 6.0 at 4 °C, followed by size exclusion using a HiLoad Superdex 75 pg 16/600 column. All ^15^N-labeled proteins were exchanged into matched buffer containing 25 mM NaPi, 150 mM NaCl, 0.5 mM DTT, pH 7.4 prior to quantification and storage as described above.

The Ub-PA activity-based probes were prepared using intein chemistry^78^, as described previously in detail^58^.

### Ub-PA reactivity assays

Ub-PA reactivity assays were performed at a 1:2, bHECT:Ub-PA molar ratio using 5 µM bHECT and 10 µM Ub-PA in reaction buffer containing 25 mM Tris, 150 mM NaCl, 0.5 mM DTT, pH 8.0. Small-scale reactions were incubated at 37 °C for 1 h. Samples were quenched with reducing Laemmli sample buffer and analyzed by SDS-PAGE.

### Gel-based E3 ligase assays

E3 ligase assays were performed using 300 nM UBA1, 2 µM Lys-less UBE2L3, 50 µM Ub (WT, K-only, K-to-R, or Y59A), with HECT E3 ligases at concentrations indicated in the figure panel or figure legend, in the presence of 5 mM ATP, 0.5 mM DTT, and 10 mM MgCl_2_. All gel-based ligase assays were performed at 37 °C. Reaction times were scaled based on the specific activity of each HECT. At the time points indicated in the figure panel or figure legend, samples were quenched with reducing Laemmli sample buffer and analyzed by SDS-PAGE.

### UbiCRest analysis

PolyUb chain assemblies using NleL, the SNc ligase, or mutants thereof, were prepared as described above. Reactions were quenched by addition of EDTA to 40 mM final concentration and DTT to 5 mM final concentration. DUBs were diluted into activation buffer containing 25 mM Tris, 150 mM NaCl, 10 mM DTT, pH 7.4 and incubated at 22 °C for 10 min, as previously described^79^. DUBs were added at 5 µM final concentration to polyUb assemblies, mixed, and incubated at 37 °C for 2 h prior to quenching in reducing Laemmli sample buffer and analysis by SDS-PAGE.

### Western blot analysis

Reactions were resolved by SDS PAGE as described above. Next, gels were transferred onto PVDF membranes using the semi-dry Trans-Blot Turbo system (BioRad) using the mixed-molecular weight setting. Following transfer, membranes were blocked at room temperature for 1 hour with TBS-T (Tris-buffered Saline with 0.1% v/v Tween-20) containing 5% milk. After blocking, membranes were washed in TBS-T. Next, membranes were incubated with an anti-Ub antibody (MilliporeSigma, MAB1510-I; 1:1,000 dilution at 4 °C overnight with gentle rocking. Membranes were again washed in TBS-T, prior to incubation with the secondary antibody (MilliporeSigma, #12-349; 1:5,000 dilution) at room temperature for 1 hour. Finally, membranes were washed again in TBS-T and then briefly incubated with Clarity ECL reagent (BioRad) and visualized by chemiluminescence scan on a Sapphire Biomolecular Imager (Azure Biosystems).

### Fluorescence-based E3 ligase (UbiReal) assays

UbiReal assays were performed as previously described^43, 44^. Fluorescence polarization (FP) was recorded using a BMG LabTech ClarioStar plate reader with an excitation wavelength of 540 nm, an LP 566 nm dichroic mirror, and an emission wavelength of 590 nm. Reactions were performed at 22 °C in low-binding Greiner 384-well small-volume HiBase microplates with 20 μL final reaction volumes.

Reactions contained 150 nM UBA1, 1 µM Lys-less UBE2L3, 37.5 µM WT (unlabeled) Ub, 10 mM MgCl_2_, 0.5 mM DTT, and NleL, SopA, or HUWE1 (or mutants thereof), at 2 µM, 2 µM, or 25 µM, respectively. Each reaction also contained 100 nM Ub with an N-terminal TAMRA fluorophore. Each reaction, in the absence of ATP, was monitored for several FP cycles, and these FP values were used as the minimum FP for the ΔFP calculation at each time point. Reactions were initiated with addition of ATP to 5 mM, and monitored over time by FP. Each reaction was performed with technical triplicates, and the average value is plotted at each time point.

### Fluorescence-based E2∼Ub discharge assays

E2∼Ub discharge assays were performed using 100 nM K6R,K48R Ub modified with an N-terminal Alexa 488 fluorophore, 300 nM UBA1, 480 nM Lys-less UBE2L3, 5 mM ATP, 5 mM MgCl_2_, and 1 mM TCEP. The mixture was allowed to react, with mixing, for 5 min at 22 °C, followed by quenching with addition of EDTA to 50 mM.

For the FP-based experiment, FP was performed as described above, but monitored using an excitation wavelength of 482 nm, an LP 504 nm dichroic mirror, and an emission wavelength of 530 nm. The reaction mixture was added to the 384-well plate and monitored over time at 22 °C. Cleavage of the E2∼Ub conjugate was initiated (time point 0 min) by addition of NleL WT or mutant to 15 nM, or addition of buffer for the negative control. FP signal was monitored over time.

For the gel-based experiment, the reaction mixture was added to tubes containing NleL WT or mutant at 15 nM final concentration, and allowed to react at 22 °C for 6 minutes. Samples were quenched with non-reducing Laemmli sample buffer, analyzed by SDS-PAGE, and visualized by fluorescence scan at 488 nm (Sapphire BioImager).

### Protein crystallization and structure determination

NleL (606-782), SopA (603-782), and VsHECT (639-847) were prepared as described above and reacted with Ub-PA at a molar ratio of 1:2 bHECT:Ub-PA overnight at 4 °C with rocking. Reactions were subsequently purified by anion exchange chromatography using a Resource Q column (Cytiva) with a 0 – 0.5 M NaCl gradient in 25mM Tris, 1 mM DTT, pH 8.5, followed by size exclusion on a HiLoad Superdex 75 pg 16/600 column (Cytiva) equilibrated with 25 mM Tris, 125 mM NaCl, 1 mM DTT, pH 7.4. NleL-Ub^D^, SopA-Ub^D^ and VsHECT-Ub^D^ were concentrated to 15 mg/mL, 9 mg/mL, and 15 mg/mL, respectively. NleL-Ub^D^ crystallized in Ligand Friendly Screen (Molecular Dimensions) in sitting drop format with 20% PEG 3350, 0.2 M KSCN, 0.1 M bis-tris propane pH 7.5, 20% glycerol, and 10% ethylene glycol at 22 °C in a 1 µL drop with 1:1 protein:precipitant ratio. SopA-Ub^D^ crystallized in hanging drop format with 22.5% PEG 8000, 0.2 M ammonium sulfate, 0.1 M sodium cacodylate pH 7.0, and 20% glycerol at 22 °C in a 1 µL drop with 1:1 protein:precipitant ratio. VsHECT-Ub^D^ crystallized in hanging drop format with 20% PEG 2K MME, 0.1 M MES pH 6.0, and 20% ethylene glycol at 22 °C in a 1 µL drop with 1:1 protein:precipitant ratio. Crystals for each bHECT-Ub^D^ were cryoprotected in mother liquor containing 25% glycerol prior to vitrification.

Diffraction data were collected at the Stanford Synchrotron Radiation Lightsource (SSRL), beamline 9-2. The data were integrated using XDS^80^ and scaled using Aimless^81^. The NleL-Ub, SopA-Ub, and VsHECT-Ub structures were determined by molecular replacement with Phaser in CCP4i2, using search models consisting of NleL (PDB: 3NB2), SopA (PDB: 2QYU), or a model of VsHECT built using Phyre2^76^, respectively, along with Ub (PDB: 1UBQ) ^13, 14, 82–84^. Automated model building was performed using ARP/wARP^85^, followed by iterative rounds of manual model building in COOT and refinement in PHENIX^86, 87^. All figures were generated using PyMOL (www.pymol.org).

### NMR analysis of NleL:UBE2D3∼Ub

The ^15^N-UBE2D3-O-^15^N-Ub conjugate was prepared using ^15^N-Ub and ^15^N-UBE2D3 C85S/S22R, as previously described^88^. NMR experiments were performed in 25 mM NaPi, 150 mM NaCl, 0.5 mM DTT, pH 7.4 with 10% D_2_O on a 500 MHz Bruker AVANCE III at 25 °C. Data were processed using NMRPipe^89^ and analyzed using NMRViewJ^90^. NMR spectra were recorded of 150 µM ^15^N UBE2D3-O-Ub alone, or following the addition of 0.1 molar equivalents (15 µM final) of NleL C753A (170-782), or 2.0 molar equivalents (300 µM final) of NleL C753A (606-782). Surface structure representations of peak broadening following NleL titration were plotted using PyMOL.

## AUTHOR CONTRIBUTIONS

TGF and JNP conceptualized the approach. TGF performed all experiments with guidance from PSB and JNP. TGF and JNP analyzed the data and wrote the manuscript with input from PSB.

## ACKNOWLEDGEMENTS

We thank David Komander (Walter and Eliza Hall Institute of Medical Research), Rachel Klevit (University of Washington), and Thomas Mund (MRC Laboratory of Molecular Biology) for sharing expression plasmids. We thank members of our laboratories and the Seattle Ub Research Group for helpful discussions. Access to NMR facilities was generously granted by Rachel Klevit, who is supported by the National Institute of General Medical Sciences (NIGMS) (1R35GM144127). Use of the Stanford Synchrotron Radiation Lightsource, SLAC National Accelerator Laboratory, is supported by the U.S. Department of Energy, Office of Science, Office of Basic Energy Sciences under Contract No. DE-AC02-76SF00515. The SSRL Structural Molecular Biology Program is supported by the DOE Office of Biological and Environmental Research, and by the National Institutes of Health, NIGMS (P30GM133894). The contents of this publication are solely the responsibility of the authors and do not necessarily represent the official views of NIGMS or NIH. This work was supported by Oregon Health & Science University (JNP), the OHSU Program in Molecular and Cellular Biosciences (5T32GM071338-14 to TGF) and the NIGMS (R35GM142486 to JNP).

## DATA AVAILABILITY

Coordinates and structure factors for the NleL-Ub^D^, SopA-Ub^D^, and VsHECT-Ub^D^ structures have been deposited in the Protein Data Bank under accession codes 8ST9, 8ST8, and 8ST7, respectively. All other data are available upon request.

