## Supplemental Files for "Bacterial mimicry of eukaryotic HECT ubiquitin ligation"

**Supplemental Figure 1: Discovery of an expanded bHECT family**

A. Percent sequence identity matrix of bHECT C-lobe regions.
B. Percent sequence identity matrix of bHECT N-lobe regions.
C. Percent sequence identity matrix of bHECT linker regions.
D. Ub ligation monitored by fluorescence polarization (FP). Reactions were initiated with ATP at timepoint 0 min. The final (75 min) ΔFP values are reported for the bHECTs NleL (4 μM), SopA (4 μM), PvHECT (4 μM), PsHECT (4 μM), EaHECT (12.5 μM), and VsHECT (4 μM).

#### Fig. S1: Discovery of an expanded bHECT family

**A**

C-lobe — % identity C-lobe

| | NleL | SopA | PvHECT | VsHECT | EaHECT | PsHECT |
|---|---|---|---|---|---|---|
| **NleL** | | 38 | 30 | 28 | 30 | 35 |
| **SopA** | 38 | | 29 | 29 | 33 | 38 |
| **PvHECT** | 30 | 29 | | 28 | 34 | 44 |
| **VsHECT** | 28 | 29 | 28 | | 21 | 31 |
| **EaHECT** | 30 | 33 | 34 | 21 | | 41 |
| **PsHECT** | 35 | 38 | 44 | 31 | 41 | |

**B**

N-lobe — % identity N-lobe

| | NleL | SopA | PvHECT | VsHECT | EaHECT | PsHECT |
|---|---|---|---|---|---|---|
| **NleL** | | 38 | 22 | 21 | 23 | 24 |
| **SopA** | 34 | | 29 | 17 | 22 | 24 |
| **PvHECT** | 22 | 19 | | 28 | 32 | 38 |
| **VsHECT** | 21 | 17 | 24 | | 8 | 25 |
| **EaHECT** | 23 | 22 | 32 | 8 | | 41 |
| **PsHECT** | 24 | 24 | 38 | 25 | 41 | |

**C**

linker — % identity linker

| | NleL | SopA | PvHECT | VsHECT | EaHECT | PsHECT |
|---|---|---|---|---|---|---|
| **NleL** | | 22 | 26 | 17 | 35 | 26 |
| **SopA** | 22 | | 4 | 26 | 4 | 17 |
| **PvHECT** | 26 | 4 | | 25 | 39 | 42 |
| **VsHECT** | 17 | 26 | 25 | | 16 | 17 |
| **EaHECT** | 35 | 4 | 39 | 16 | | 30 |
| **PsHECT** | 26 | 17 | 42 | 17 | 30 | |

**D**

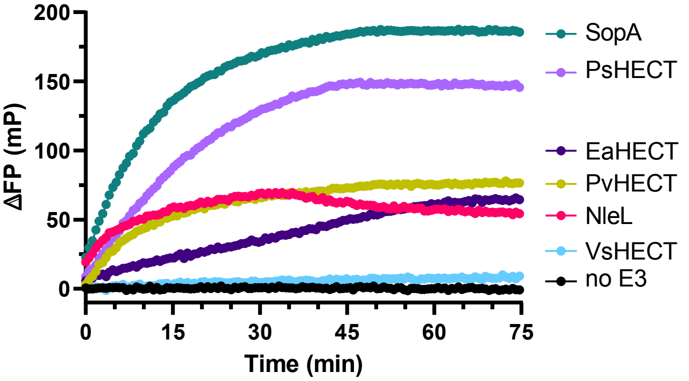


| | Δ FP (75 min.) |
|---|---|
| SopA | 186 |
| PsHECT | 146 |
| EaHECT | 76.6 |
| PvHECT | 64.4 |
| NleL | 54.4 |
| VsHECT | 9.20 |

**Supplemental Figure 2: Structural and biochemical analysis of bHECT C-lobes**

A. Full 2|Fo-Fc| electron density for the NleL-$Ub^D$ structure, shown at 1σ.
B. Full 2|Fo-Fc| electron density for the SopA-$Ub^D$ structure, shown at 1σ.
C. Full 2|Fo-Fc| electron density for the VsHECT-$Ub^D$ structure, shown at 1σ.
D. Overlay of the NleL-$Ub^D$ structure with two apo NleL structures (PDB: 3NB2 and 3NAW), aligned on their C-lobes. NleL β-helix, N-lobe, and C-lobe domains are labeled.
E. Overlay of the SopA-$Ub^D$ structure with two apo SopA structures (PDB: 2QZA and 2QYU), aligned on their C-lobes. SopA β-helix, N-lobe, and C-lobe domains are labeled.

### Fig. S2: Biochemical and structural analysis of bHECT C-lobes

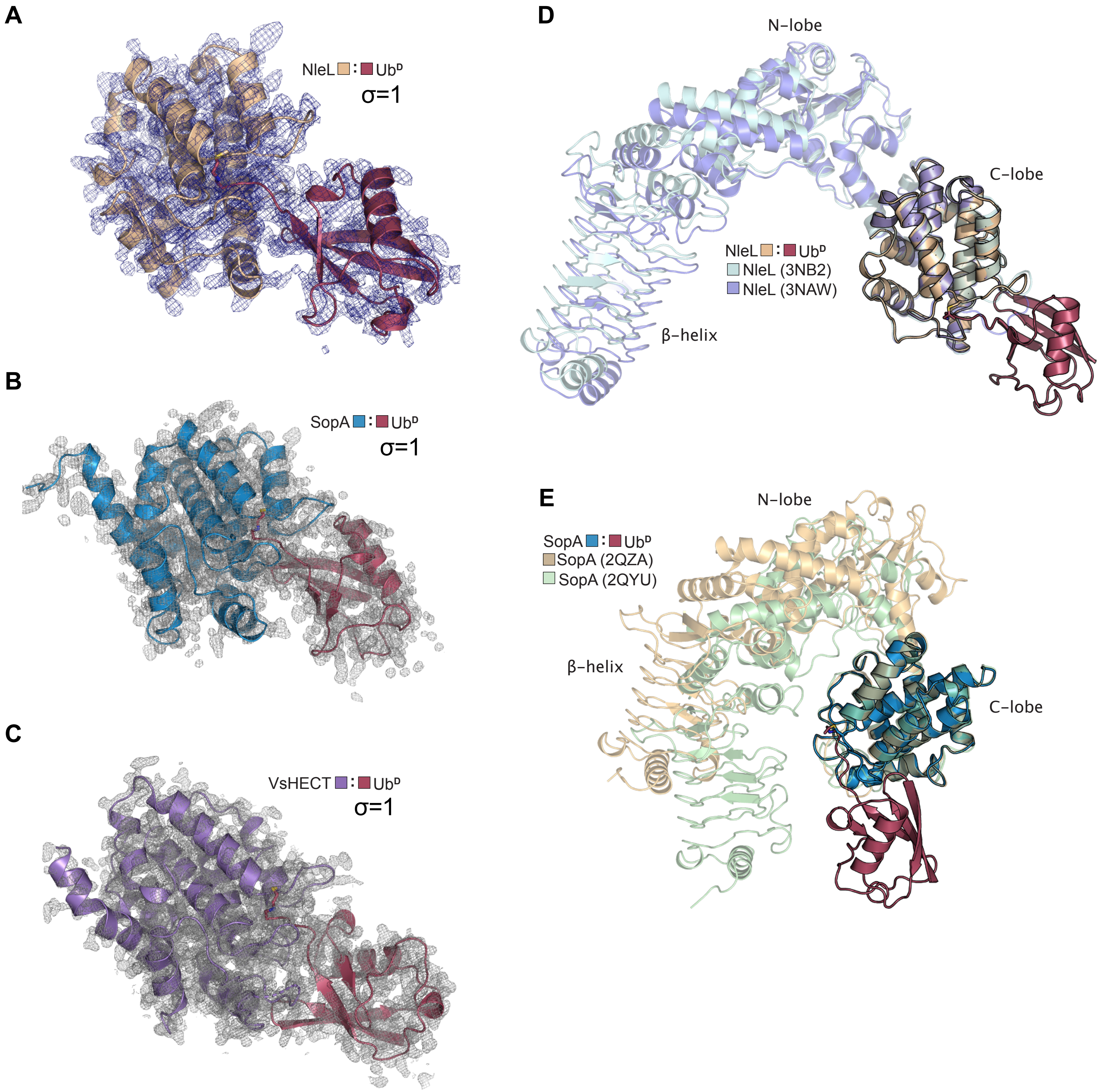

**Supplemental Figure 3: bHECT activation of Ub$^{D}$**

A. Structures of SopA-Ub$^{D}$, NleL-Ub$^{D}$, and VsHECT-Ub$^{D}$, focusing on the hydrogen bonding networks established between the C-lobe and Ub$^{D}$. Hydrogen bonds are shown with black dashes.

B. Multiple sequence alignment (MSA) of the bHECTs, focused on the C-lobe. The sequences of the acidic loops and Cys loops are highlighted. Asterisks and red boxes are used to indicated the conserved Ub$^{D}$-coordinating Arg residue, the acidic loop Glu residue, the partially-conserved Cys loop Phe residue, and the conserved active site Cys.

C. Gel-based Ub-PA reactivity assay with WT or the Ub$^{D}$-coordinating Arg-to-Ala mutant for the full HECT (FH) domain NleL, FH domain SopA, and C-lobe (C-l) of VsHECT. 5 μM of bHECTs were reacted with 10 μM Ub-PA for 2 hr at 22 °C. Reactions were quenched and resolved by SDS-PAGE with Coomassie staining.

D. Structure of SopA-Ub$^{D}$, highlighting the unique interactions observed at the C-lobe:Ub$^{D}$ interface.

E. Structure of VsHECT-Ub$^{D}$, highlighting the unique interactions observed at the C-lobe:Ub$^{D}$ interface.

F. MSA of the bHECTs, highlighting the sequence insertion that partially mediates the unique Ub$^{D}$ interactions observed in the VsHECT C-lobe shown in **E**. VsHECT residues I677, I712, and Y713 from **E** are marked with red asterisks.

G. Gel-based Ub ligase assay for WT VsHECT and structure-guided mutants as monitored by formation of diUb. Reactions were quenched, resolved by SDS-PAGE, and visualized by Western blot for Ub.

#### Fig. S3: bHECT activation of donor Ub

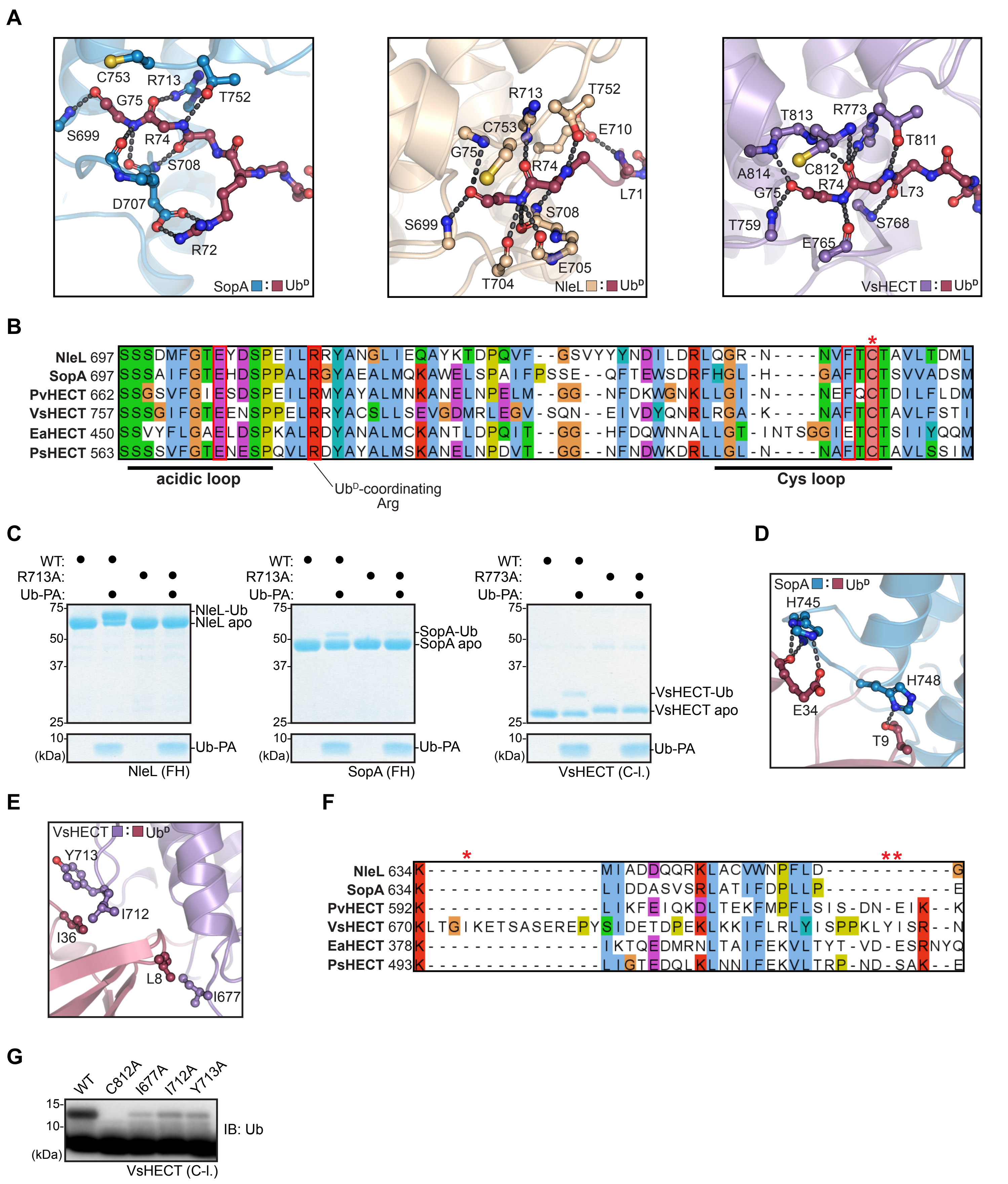

**Supplemental Figure 4: Model for E2-bHECT transthiolation**

A. The NleL-Ub$^{D}$ and NleL:UBE2L3 (PDB: 3SQV) overlay, representing a model of the E2:NleL~Ub intermediate, overlayed with a UBE2D3 structure (PDB: 3UGB) to highlight the similarity when using either E2. UBE2D3 was aligned onto UBE2L3 (C$\alpha$ RMSD=1.006Å). The E2:Cys loop and E2:N-lobe interfaces are highlighted, with the conserved Phe residues mediating the interaction between E2 and NleL N-lobe shown as sticks, and the active site Cys residues for both NleL and UBE2L3/UBE2D3 shown as yellow spheres.

B. 2|Fo-Fc| electron density for the entire Cys loop in the NleL-Ub$^{D}$ structure at 1$\sigma$. Residues on the Cys loop are indicated, as well as the $\alpha$-helix numbers.

C. As in **B**, for the SopA-Ub$^{D}$ structure

D. Histogram of relative peak intensities for UBE2D3 and Ub resonances from $^{1}$H,$^{15}$N-TROSY spectra of $^{15}$N-labeled UBE2D3-O-Ub conjugate, following titration of 0.1 molar equivalencies (eqv) of the full NleL HECT-like domain containing the catalytic C753A mutation. Peak intensities are relative to $^{15}$N-labeled UBE2D3-O-Ub conjugate alone. Grey bars indicate unassigned or Pro residues. Yellow bars indicate relative intensity changes greater than one standard deviation from the mean.

E. Histogram of relative peak intensities for UBE2D3 and Ub resonances from $^{1}$H,$^{15}$N-TROSY spectra of $^{15}$N-labeled UBE2D3-O-Ub conjugate, following titration of 2.0 molar equivalencies (eqv) of NleL C-lobe C753A. Peak intensities are relative to $^{15}$N-labeled UBE2D3-O-Ub conjugate alone. Grey bars indicate unassigned or Pro residues. Yellow bars indicate relative intensity changes greater than one standard deviation from the mean.

F. Validation of the E2~Ub FP discharge assay. FP values of Alexa 488-labeled Ub K6R, K48R change relative to molecular weight as E1 is added to form E1~Ub, UBE3L3$^{K0}$ is added to form UBE3L3$^{K0}$~Ub, and SopA is added resulting in E2~Ub discharge to free Ub via a transient E3~Ub intermediate.

#### Fig. S4: Model for E2-bHECT transthiolation

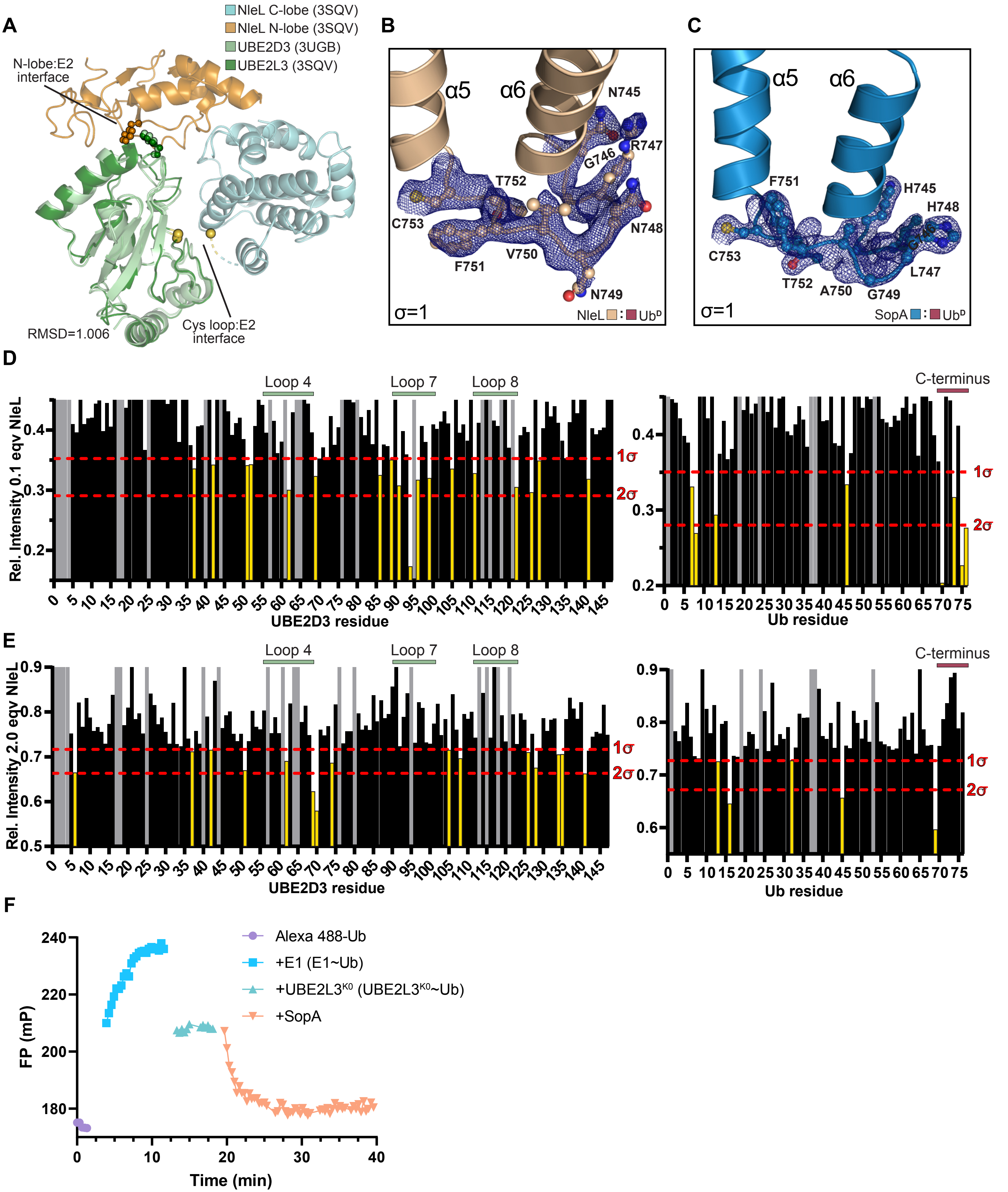

**Supplemental Figure 5: bHECT coordination of Ub$^{A}$**

A. 2|Fo-Fc| electron density for key interactions at the Ub$^{A}$:NleL-Ub$^{D}$ interface, shown at 1σ. Acidic loop residue E705, Cys loop residues C753 and F751, and Ub$^{A}$ residues K48 and Y59 are shown.
B. Gel-based polyUb specificity assay for the NleL C-lobe, using the panel of K-only Ub mutants. Reactions were quenched and resolved by SDS-PAGE with Coomassie staining.
C. As in **B**, for the SopA C-lobe.
D. Schematics of the HECT-like domains of SopA and NleL, and the C-lobe domain swap used to generate the SopA-NleL chimera (SNc) and SNc E705A constructs. PolyUb specificity of each construct is indicated.
E. Gel-based polyUb specificity assay using the panel of K-only Ub mutants for the SNc chimera without (left) and with (right) the E705A acidic loop mutation. Reactions were quenched and resolved by SDS-PAGE with Coomassie staining.

#### Fig. S5: bHECT coordination of acceptor Ub

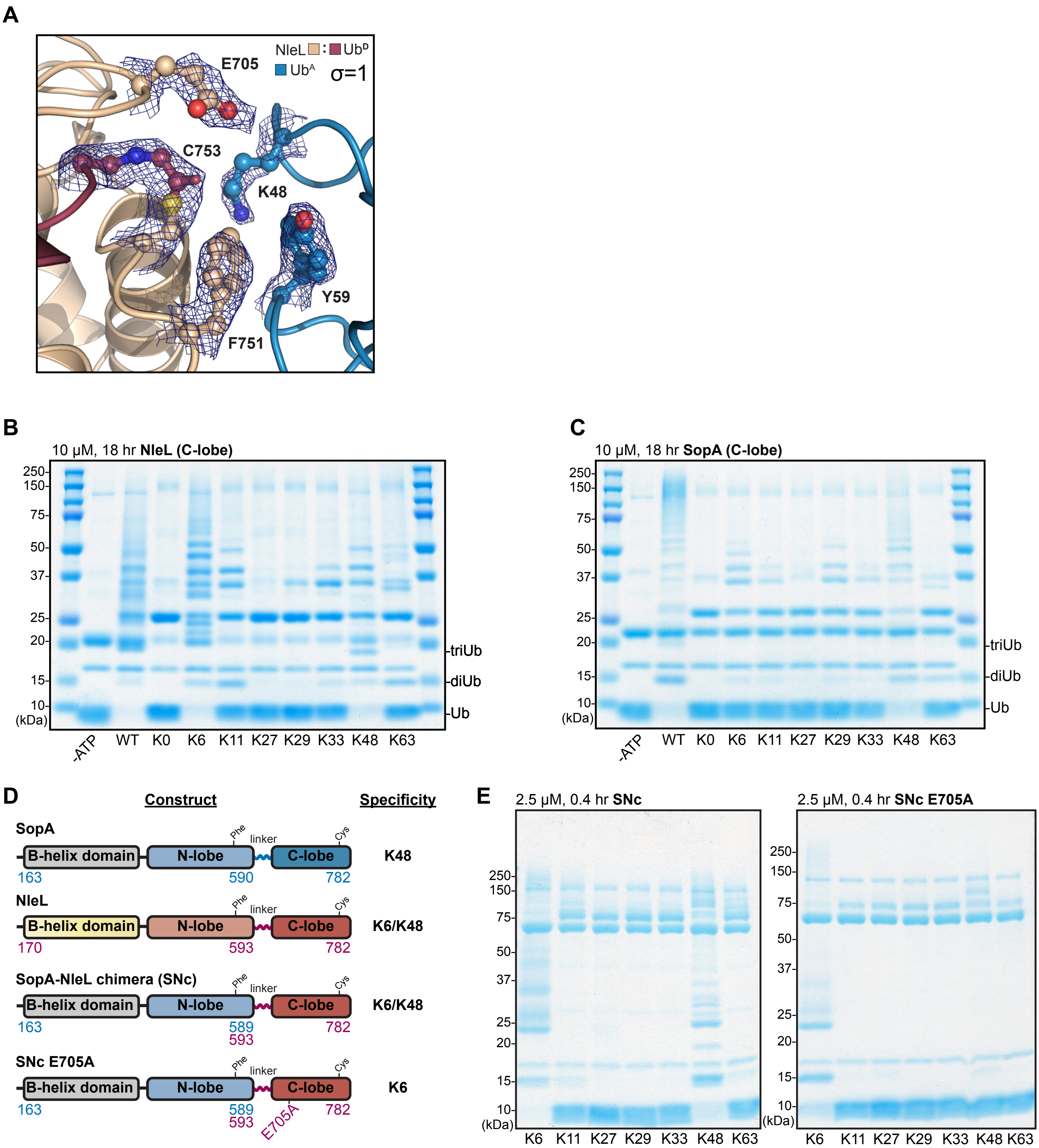

**Supplemental Figure 6: HUWE1 acidic loop mutants show increased K6 polyUb ligation**

A. MSA of selected eHECTs, highlighting the region of the C-lobe surrounding the active site. Asterisks and red boxes are used to indicate the conserved active site Cys and the conserved Cys-loop Phe. The sequences of the Cys loop and potential acidic loop are highlighted. Red boxes indicate potential acidic residues.

B. Overlay of the HUWE1-Ub$^{D}$ (PDB: 6XZ1) structure with Ub$^{A}$ from the Ub$^{A}$:NleL-Ub$^{D}$ structure, showing the Cys loop and C-lobe acidic loop of HUWE1, along with the Ub$^{A}$ residues important for NleL K48-linked Ub ligation, Y59 and K48.

C. Structure of apo HUWE1 (PDB: 5LP8), focusing on the active site, with the C-lobe shown in teal and the N-lobe shown in gold. The C-lobe acidic loop containing E4315 is shown, as well as a second acidic loop from the T conformation of the N-lobe. Sequence conservation of the second N-lobe acidic loop is shown with other eHECTs, with the location of acidic residue selected for mutational analysis indicated with a blue star and blue box.

D. View of the HUWE1-Ub$^{D}$ structure (PDB: 6XZ1), overlayed with the N-lobe of Rsp5 (PDB: 4LCD). The acidic loop region lacking electron density in the Rsp5 structure is shown with the dashed line. Acidic loop and Cys loop residues for HUWE1 are shown.

E. Gel-based polyUb specificity assay for HUWE1 WT and the N-lobe acidic loop mutants, using a subset of K-only Ub mutants. Reactions were quenched and resolved by SDS-PAGE with Coomassie staining. Gel regions corresponding to monoUb, diUb, and triUb are shown for clarity.

### Fig. S6: HUWE1 acidic loop mutants show increased K6 polyUb ligation

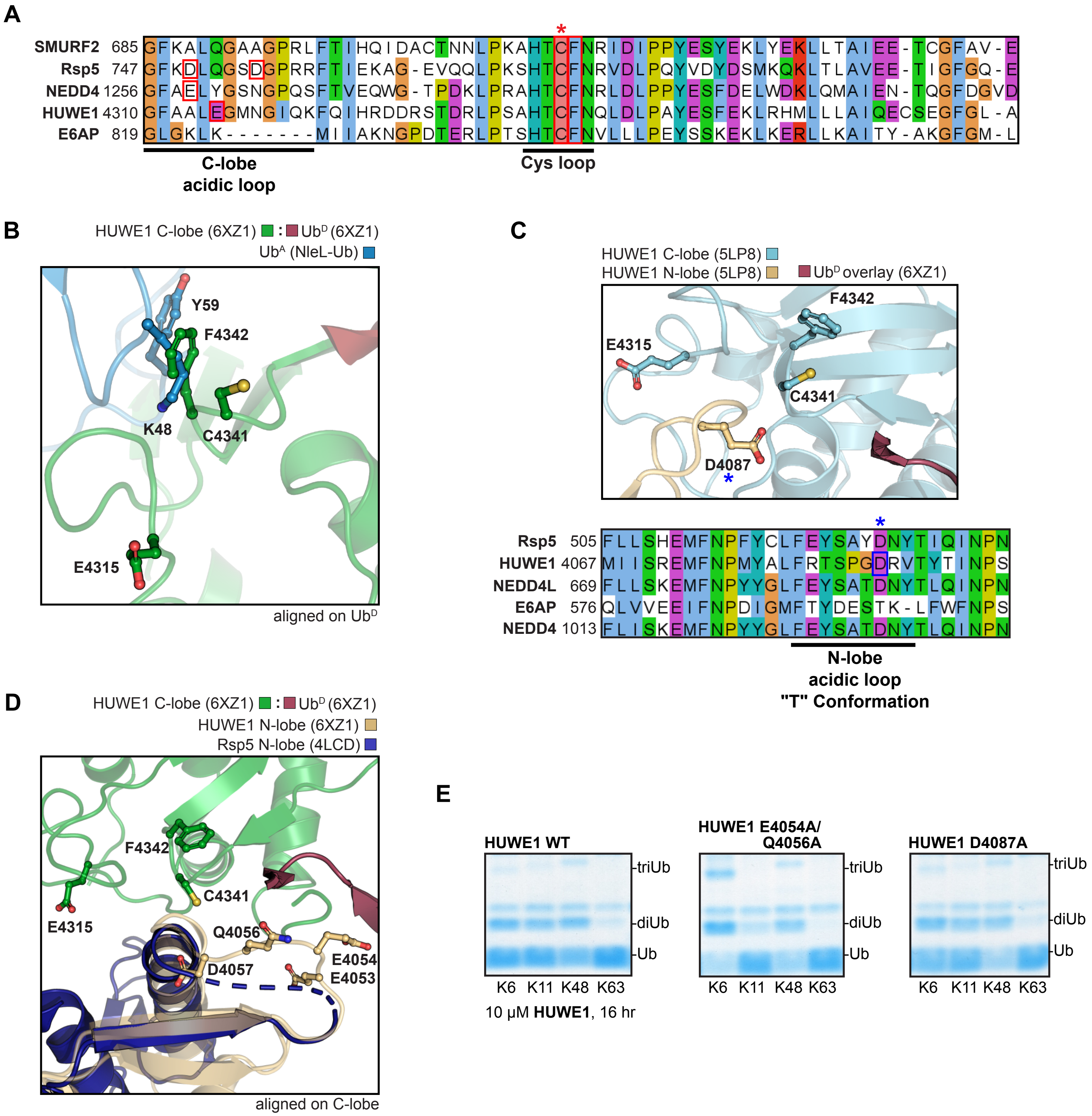

**Supplementary Table 1: Constructs used in this study**

| Gene name | Organism | Uniprot ID | NCBI/GenBank | Construct |
|---|---|---|---|---|
| *nleL* | Enterrohemmorhagic *Escherichia coli* O157:H7 str. Sakai | A0A0H3JDV8 | WP_001301673.1 | 170-782 (FH)<br>606-782 (C-lobe) |
| *sopA* | *Salmonella enterica* Typhimurium str. SL1344 | Q8ZNR3 | WP_000703998.1 | 342-782 (FH)<br>603-782 (C-lobe) |
| *vsHECT* | *Verrucomicrobia species* | A0A2V2RSR1 | PWU08673.1 | 639-847 (C-lobe) |
| *pvHECT* | *Proteus vulgaris* | A0A292CDM3 | WP_192940890.1 | 292-745 (FH) |
| *psHECT* | *Pantoea stewartii* | H3RGA3 | WP_006120546.1 | 210-646 (FH)<br>464-646 (C-lobe) |
| *eaHECT* | *Erwinia amylorova* | D4HXM4 | WP_013036135.1 | 85-537 (FH) |
